# Chemokine-binding all-D-CLIPS™ peptides identified using mirror-image phage display

**DOI:** 10.1101/2025.09.09.675118

**Authors:** Stepan S. Denisov, Emilia L. Bialek, Fabio Beretta, Gintare Smagurauskaite, Johannes H. Ippel, Eline Fijlstra, Sangram S. Kale, Peter Timmerman, Tilman M. Hackeng, Paul Proost, Michael Goldflam, Ingrid Dijkgraaf

**Affiliations:** Insitute of Biological Chemistry, University of Vienna, Währinger Str. 38, 1090 Vienna, Austria; Department of Biochemistry, Maastricht University, Cardiovascular Research Institute Maastricht (CARIM), Universiteitssingel 50, 6229 ER, Maastricht, the Netherlands; Rega Institute, KU Leuven, Herestraat 49, 3000, Leuven, Belgium; RDM Cardiovascular Medicine, University of Oxford, Roosevelt Drive, Oxford OX3 7BN, UK; Biosynth B.V., Zuidersluisweg 2, 8243 RC Lelystad, the Netherlands

**Author notes:** Authors contributed equally (Michael Goldflam, Ingrid Dijkgraaf).

## Abstract

Chemokines are secreted blood proteins, which steer leukocyte migration in the inflammatory response. Neutralization of chemokines is believed to be a beneficial therapeutic strategy for the treatment of inflammation-associated diseases. Proteolytically stable chemokine-binding peptides could be suitable candidates for the development of chemokine-neutralizing agents. Here, we report mirror-image phage display selection of cyclic all-D-peptides against the C-X-C motif chemokine ligand 8 (CXCL8). Selection yielded structurally diverse all-D-peptides with sub-micromolar affinity to the target CXCL8 chemokine and different selectivity to related chemokines. Binding of these all-D-peptides caused dissociation of the native CXCL8 dimer and disruption of its binding to GAGs, without effect on in vitro cell migration. This work demonstrates the example of mirror-image phage display selection of cyclized all-D-peptides and its utility for the development of chemokine-binding agents.

## Introduction

Chemokines (chemotactic cytokines) are small secreted proteins that orchestrate cell trafficking by activating G protein coupled chemokine receptors^1^. Since they play a crucial role in the progression of the inflammatory response, chemokine neutralization could provide a promising strategy for the treatment of inflammation-associated conditions, such as atherosclerosis^2^, myocardial infarction^3^, stroke^4^, and arthritis^5^. Due to a relatively smooth surface, neutralization of chemokines by small molecules is challenging with only several examples focused on CCL2^6,7^ and CXCL12^8–10^ described to date. Peptides, with molecular weight 500 – 5000 Da, represent a vast chemical space for selection of binders against undruggable targets^11^. Taking into account the extracellular nature of chemokines and, therefore, no need to overcome peptides’ intrinsic low membrane permeability^12^, they draw attention as prospective targets for chemokine-binding agents. Although many examples of peptides, either rationally designed^13^, selected from combinatorial libraries^14^, derived from chemokine receptors,^15^ or based on tick chemokine-binding proteins^16–18^ peptides are reported, poor metabolic stability of peptides greatly hampers their therapeutic application. Application of proteolytic resistant peptidomimetics, such as N-substituted oligoglycines e.g. peptoids, was reported for the development of CXCL8-neutralizing agents^19,20^. The use of D-amino acids instead of proteinogenic L-enantiomers is another highly effective way to protect a peptide bond from proteolytic cleavage^21^. Since high proteolytic stability also increases peptide oral bioavailability^22^ and decreases immunogenicity^23,24^, the use of D-peptides could be a superior approach for chemokine-binding peptide development.

All-D-peptides can be selected using the ‘mirror-image’ display approach^25,26^, which relies on the search for binding sequences in the chemical space of natural L-peptides against a non-natural D-protein target. Subsequent synthesis of selected sequences in D-enantiomeric form yields the non-natural variant of those binders for the corresponding natural L-protein target. Using the natural biomolecular machinery, this approach allows for the encoding and screening of billions of peptide sequences, which surpasses the capacity of any synthetic library of peptidomimetics^27,28^. Mirror-image mRNA display was recently used to select high-affinity cyclic sulphotyrosine-containing D-peptides against CCL22^29^ and D-monobodies against CCL2^30^. Being compatible with the chemical linkage of peptides onto scaffolds (CLIPS™) technology^31,32^, which yields structurally constrained cyclic peptides and increases their affinity, we envisioned that using the mirror-image phage display could be used for the development of cyclic chemokine-neutralizing all-D-peptides. Here, we report the development of cyclic all-D-peptides against C-X-C motif chemokine ligand 8 (CXCL8, interleukin-8, IL-8), which plays a crucial role in neutrophil recruitment and/or the strengthening of angiogenic responses during development of inflammation and progression of colorectal cancer, HIV-associated neurocognitive disorder, and psoriasis^33–36^.

## Results and discussion

### Synthesis of D-CXCL8

Despite impressive progress in creating mirror-image replication, transcription, and translation systems^37^, chemical synthesis remains the only method to obtain all-D-proteins. To allow for mirror-image selection, the biotinylated D-amino acid variant of CXCL8 (IL-8(6-77) UniprotID p10145) was assembled from three synthetic fragments (Fig. S1-3) using native chemical ligation (NCL)^38^ (Fig. 1A). Biotin was conjugated via a C-terminal Lys(Gly-Gly) linker to fragment 2 to allow immobilization of D-CXCL8 on magnetic beads. After NCL of fragments 1 and 2 (Fig. S4), the N-terminal thiaproline residue was deprotected with *N*-methylhydroxylamine, yielding fragment 4 that was subsequently ligated to fragment 3 (Fig. S5). After completion of the second NCL reaction, full-length protein 5 was refolded under oxidative conditions, resulting in D-CXCL8-biotin 6 (Fig. S6). LC-MS analysis of 6 (Fig. 1B-C) showed the presence of a single isomer and a 4 Da shift in molecular weight compared to 5, indicating the formation of the two disulfide bonds. CD spectrum of D-CXCL-biotin mirrored the spectrum of L-CXCL8-biotin, thus proving the correct folding of the obtained D-protein (Fig.1D).

**Figure 1.**
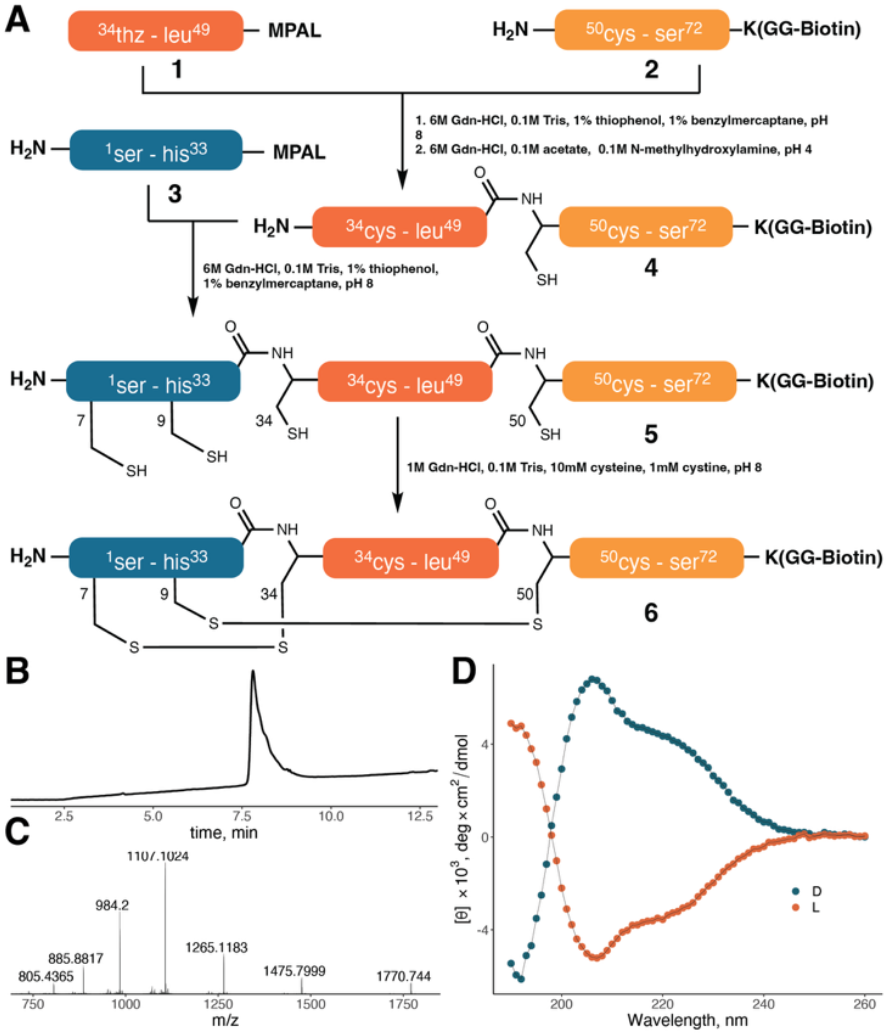
**A**. Synthetic route to D-CXCL8-biotin. HPLC trace (**B**) and ESI mass spectrum (**C**) of D-CXCL8-biotin (**6**). Observed deconvoluted [M+H]^+^ is 8844.8 Da, calculated [M+H]^+^-8844.8 Da. **D**. CD spectra of D-(blue) and L-(orange) CXCL8-biotin.

### Mirror-image phage display selection

For the selection procedure, a phage library of peptides with a randomized region of 10 amino acids was used. The diversity of the library was estimated to be 3.4×10^9^ based on transformation efficiency. As cyclization is known to increase the binding affinity by limiting conformational flexibility, the randomized region was flanked by Ala-Cys and Cys-Gly from the N- and C-terminus, respectively, to enable further cyclization with 1,3-bis(bromomethyl)benzene (T2 scaffold)^31^ (Fig. 2A). The phage library was panned against D-CXCL8-biotin immobilized on streptavidin beads for three rounds of phage selection. The recovery rates of 4×10^-7^, 2×10^-4^, and 8×10^-4^ were observed for consecutive selection rounds, representing a ∼2000-fold enrichment. After the third round of selection, the resulting phage population was subjected to the next-generation sequencing (NGS). Approximately ∼17% of sequences in the obtained pool contained one of the streptavidin-binding motifs^39^ and were filtered out. To analyze the chemical space of selected peptides, a pair-wise sequence dissimilarity matrix was calculated for the variable region of the 500 most abundant peptides and projected on a 2D plane using metric multidimensional scaling (MMDS). This revealed the presence of 4 clusters with 62, 101, 276, and 61 sequences, respectively (Fig. 2B). pLogo plots^40^ were used to identify conserved sequence motifs within the clusters (Fig. 2C). Cluster 1 and 4 display the Ile-(Xxx)_2_-Asp-(Xxx)_2_-Glu-Tyr motif shifted by one position to each other. In cluster 2, a similar motif with hydrophobic and aromatic residues separated by a stretch of acidic amino acids was observed with Trp instead of Tyr in the C-terminus. No prevalent sequence motif was found in cluster 3.

**Figure 2.**
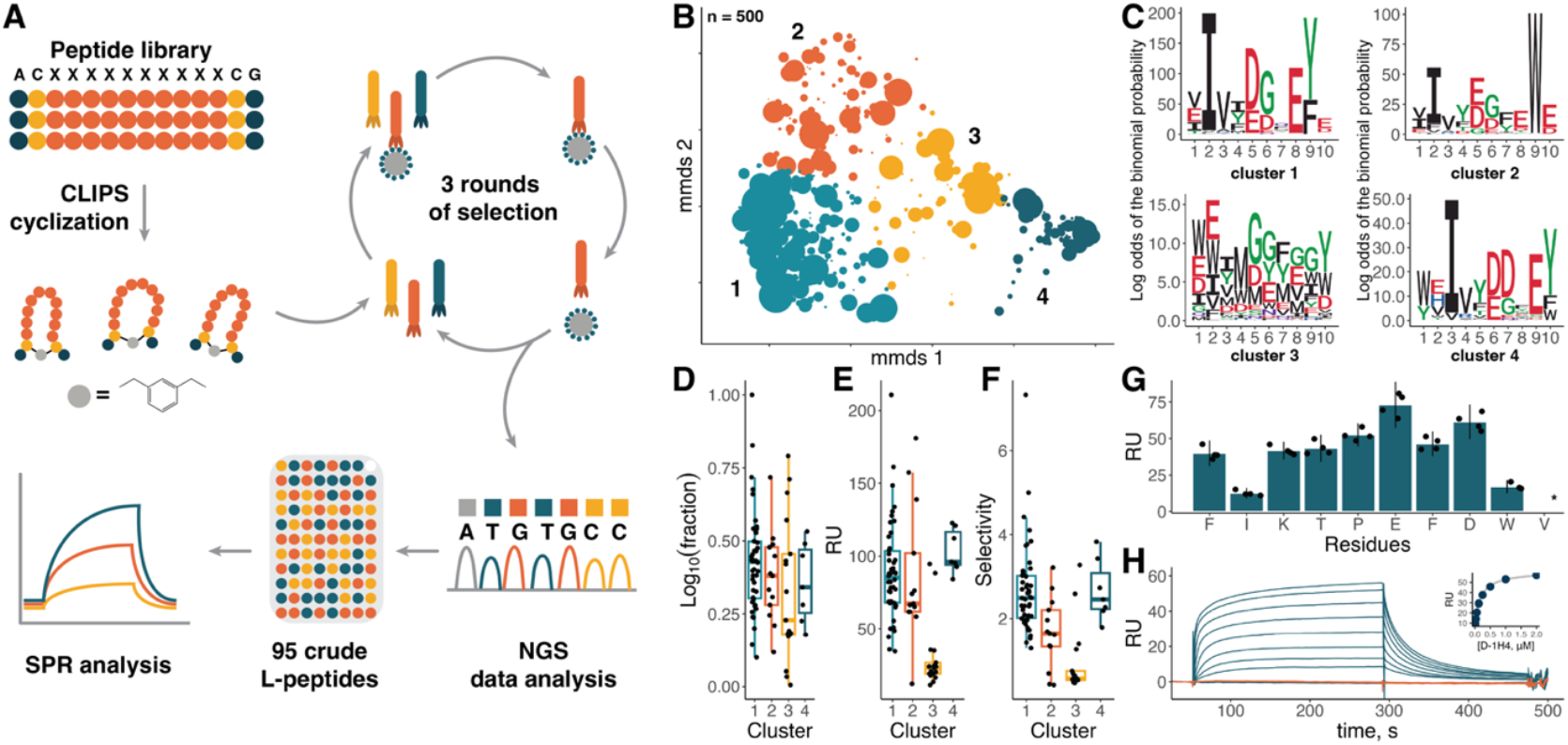
**A**. Scheme illustrating the phage display selection pipeline. **B**. 2D projection of the chemical space of 500 of the most abundant sequences after phage display selection based on sequence dissimilarity. The markers’ size is scaled proportionally to log_10_ of abundance in the NGS pool, clusters are shown in color and numbered. **C**. pLogo plots for four clusters indicated in B. Abundancy (**D**), binding level to D-CXCL8 (**E**), and selectivity (**F**) of crude L-peptides selected for screening. Abundancy is expressed as log_10_ of peptides’ fraction in the NGS pool, selectivity as a ratio of binding levels to D- and L-CXCL8. **G**. Binding level of unpurified Ala-mutants of L-1H4 peptide to D-CXCL8. **H**. Interaction sensograms of D-1H4 with L-CXCL8 (blue) and D-CXCL8 (orange) at concentrations ranging from 15 nM to 2 μM. The binding curve based on equilibrium values is shown in the insert.

### Library screening and lead peptide characterization

For further screening, 95 sequences (9, 18, 50, and 18 from clusters 1, 2, 3, and 4, respectively) covering different levels of abundance (Fig. 2D) were synthesized by Fmoc-based SPPS and cyclized by a CLIPS™ T2-scaffold. This crude CLIPS™ L-peptide library was used to test the binding to D-CXCL8 using surface plasmon resonance (SPR) biosensor analysis. Whereas 12 peptides showed high non-specific binding to the chip surface and were discarded, binding of the remaining peptides was tested at 10 μM using the biosensor chip with ∼3 kRU of immobilised D-CXCL8-biotin (Fig. 2E, S7). Peptides from clusters 1, 2, and 4 showed varying levels of binding with mean values of 90, 86, and 103 RU, respectively. In contrast, peptides from cluster 3 demonstrated a significantly lower binding, only reaching 30 RU on average, which could be attributed to the bulk effect caused by uncompensated impurities from the crude synthetic mixture during the SPR experiment. Taken together with the lack of a conserved motif, this could indicate that cluster 3 mainly consists of non-specific binders. To investigate the importance of a conserved motif found within clusters, alanine scanning of the L-1H4 peptide from cluster 2 was performed (Fig. 2G). Substitution of Ile2 and Trp9, which are highly prevalent in cluster 2, led to a substantial drop in the binding level compared to less conserved residues. This exemplifies the importance of sequence clustering for improving the hit peptide selection. Additionally, the binding of crude non-cyclized L-1H4 to L- and D-CXCL8 was tested. The linear peptide showed only non-specific binding (data not shown), indicating the essential role of CLIPS™-cyclization. To ensure the stereospecificity of the binding peptides, binding to L-CXCL8 was additionally measured (Fig. 2F). Peptides from clusters 1 and 4 demonstrated a mean selectivity of 2.7, whereas for cluster 2 the value was lower and reached 1.7. Eventually, 4 peptides from 3 different clusters - 1H4, 2A5, 3A11, and 3A12 were synthesized and purified as CLIPS™ L- and D-peptides (Fig. S8-15). When tested in the SPR biosensor assay, CLIPS™ D-peptides bound L-CXCL8, but not D-CXCL8, in a dose-dependent manner (Fig. 2H, S16) with sub-µM apparent K_D_ values (Table 1).

**Table 1.**
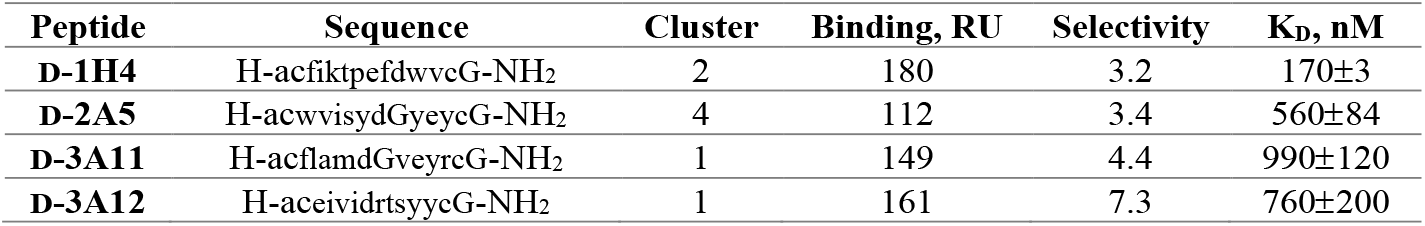
The summary table of selected D-peptides. K_D_ values are calculated as a mean of three technical replicates.

Since chemokines compose a large family of structurally conserved proteins, both nature-derived and selected from combinatorial libraries chemokine-binding peptides often bind multiple chemokines^14,17^. To assess the selectivity of the CLIPS™ D-peptides, their binding to L-CXCL1 (42% identity with L-CXCL8), L-CXCL4 (35%), and L-CCL5 (27%) was assessed by SPR analysis (Fig. 3A). D-3A11, D-3A12, and D-1H4 showed no significant binding to all tested chemokines at a concentration of 2 µM. In contrast, D-2A5 bound both L-CXCL4 and L-CCL5, but not L-CXCL1, in a dose-dependent manner (Fig. S17). Tyrosine and sulfotyrosine residues are known to be crucial for binding the N-terminus of chemokine receptors CCR5 and CXCR3 to CCL5 and CXCL4, respectively^41–43^. A similar role was shown for Tyr in class A evasins – CC-type chemokine-binding proteins from the tick saliva^44^. Thus, the presence of three Tyr in D-2A5 could be a possible explanation for the off-target binding to CCL5 and CXCL4. Given that L-CCL5 and the L-CCL5/L-CXCL4 heterodimer are considered promising targets for the atherosclerosis treatment^2^, this serendipitous result could be beneficial for the development of CLIPS™ D-peptides against atherosclerotic inflammation.

**Figure 3.**
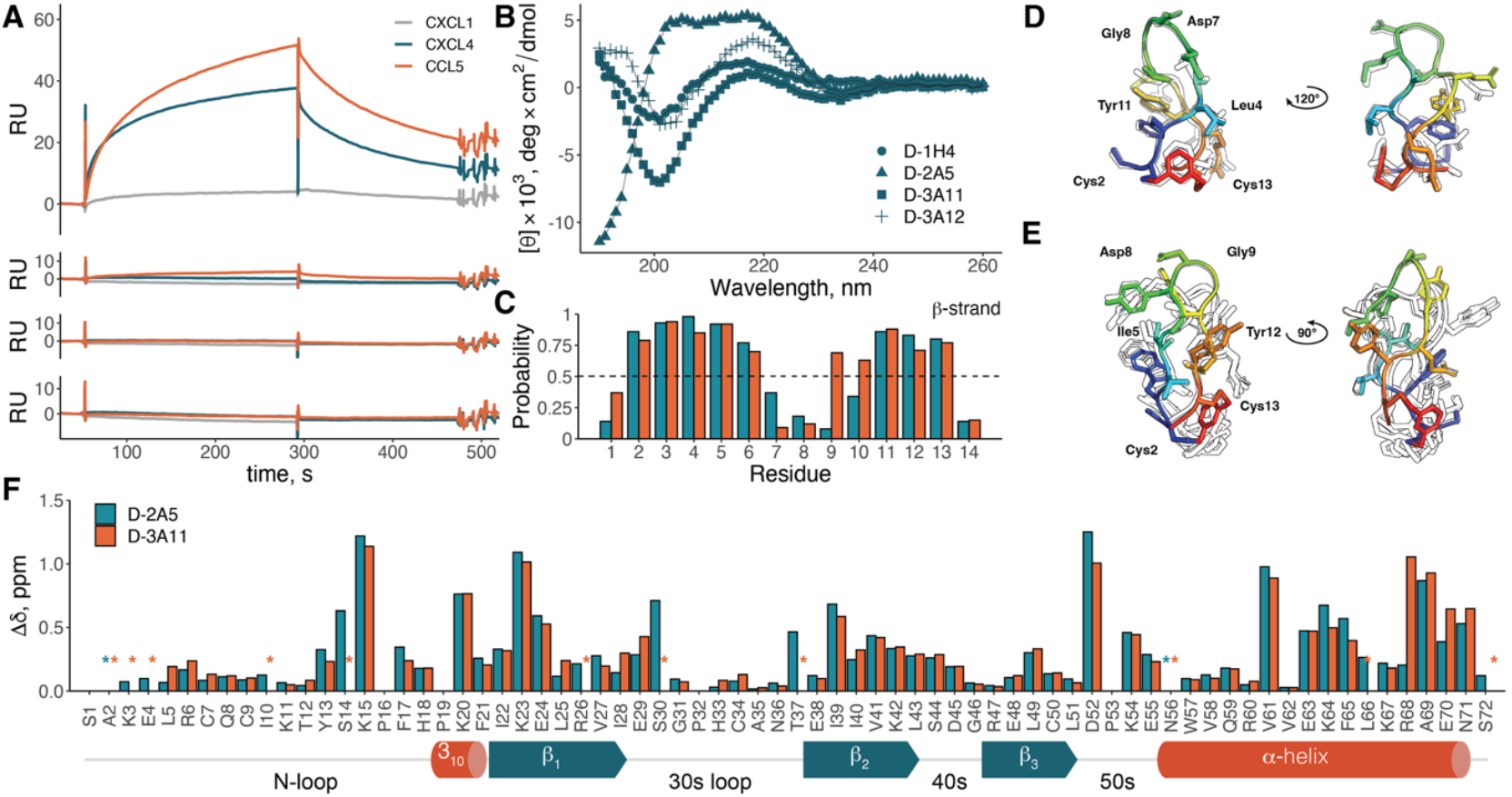
**A**. Sensograms of the interaction of 2 µM of CLIPS™ D-peptides with L-CXCL1, L-CXCL4, and L-CCL5. B. CD spectra of 0.1 mg/mL all-D-CLIPS™ peptides. C. CSI secondary structure prediction for D-3A11 in water (blue) and D-2A5 in DMSO (orange). Only the β-strand probability is shown for visibility. Overlay of 10 lowest energy models calculated for D-3A11 in water (**D**) and D-2A5 in DMSO (**E**). The top model is shown in colour. **F**. Weighted ^15^N-^1^H CSP plot for [^15^N, ^13^C] L-CXCL8 dimer and L-CXCL8 complex with D-2A5 (blue) and D-3A11 (orange). Missing signals are indicated by ^*^. The schematic representation of the CXCL8 dimer secondary structure is shown at the bottom.

We next assessed whether such a difference in selectivity is reflected in peptides’ structure using CD and NMR spectroscopy. CLIPS™ L- and D-peptides have mirrored CD spectra with a distinct pattern of signals (Fig. 3B, S18). Spectra of L-1H4, L-3A11, and L-3A12 showed positive peaks at 200 nm and 230 nm and a negative peak at 220 nm, which was previously observed for CLIPS-peptides and attributed to an anti-parallel β-sheet^32^. In contrast, the CD spectrum of L-2A5 has a broad negative peak at 203-220 nm with a strong positive one at 190 nm. However, prediction by the BeStSel algorithm^45^ indicated that all four peptides contain anti-parallel β-strand and turn secondary structure elements (Fig, S19), which could be related to the high variability of CD spectra for β-strand structures^46^.

We next employed NMR spectroscopy to further study the structure of selected D-peptides. Unfortunately, the peptides’ low solubility and aggregation at high concentrations strongly hampered the analysis, and therefore, it was only possible to obtain the full assignment of the observed signals in an aqueous solution for D-3A11 (Table S1). Analysis of D-3A11 using the chemical shift index (CSI)^47^ for Cα, Cβ, Hα, and NH atoms showed the presence of two β-strands in regions D-Cys2-D-Met6 and D-Val9-D-Cys13 separated by the Asp-Gly motif (Fig. 3C). Calculation of the structure for D-3A11 was performed by simulated annealing with the eefx^48,49^ force field using 258 unambiguously assigned inter-residue ^1^H-^1^H NOE signals and 16 chemical shift derived dihedral angles as distance and angle restraints, respectively. D-3A11 forms a tight twisted hairpin with an Asp-Gly turn on the tip and the T2-CLIPS™ scaffold at the base of the loop (Fig. 3D). D-Phe3 and D-Tyr11 are involved in π-π stacking, which causes distortion of the torsion angles (Fig. S20). In the case of D-2A5, a sufficient signal level and unambiguous assignment were only achieved in DMSO (Table S1). Although DMSO is known to destabilize protein and peptide structures, CSI analysis of D-2A5 showed the presence of two β-strands (Fig. 3C). In contrast to D-3A11, β-strands are separated by a stretch of four unstructured residues D-Tyr7-D-Tyr10. The structure of D-2A5 was calculated using 146 intra-residue distance and 19 dihedral restraints (Fig. 3F). Compared to D-3A11, D-2A5 showed a much higher structural plasticity as evidenced by 0.52 and 2.1 Å values of all-atom RMSD for the 10 lowest-energy models, respectively. The crucial difference between D-2A5 and D-3A11 is the relative position of the T2-CLIPS™ scaffold. In D-2A5 the dimethyl-benzene moiety is turned almost 90° compared to D-3A11, which creates an overhang at the N-terminus, contrasting to D-3A11 where the hairpin adopts a symmetrical shape. This distortion also brings Ile5 and Tyr12 in D-2A5 close to each other, similarly to Leu4 and Tyr11 in D-3A11. Taking into account that both peptides share an Ile/Leu-(Xxx)2-Asp-Gly-(Xxx)2-Tyr motif, that indicates that spatial organization of residues from the conservative pattern detected in NGS data is important for binding and was a selection criterion during phage display selection. To shed light on this mechanism of binding, metabolically enriched [^15^N, ^13^C] L-CXCL8 was studied using NMR spectroscopy upon binding D-2A5 and D-3A11. At μM concentrations used in NMR experiments, one set of signals was observed in the ^15^N-^1^H HSQC spectrum of L-CXCL8 attributed to a homodimer^50^ (Fig. S21). Addition of 300 μM of D-2A5 to 59 μM [^15^N, ^13^C] L-CXCL8 resulted in a new set of amide signals, from which all but Ala2 and Asn56 were assigned (Fig. S22). In the case of D-3A11, addition of 300 μM of peptide to 75 μM of [^15^N, ^13^C] L-CXCL8 did not lead to the full formation of the complex, yielding two sets of amide signals, one for the CXCL8-dimer and another for the complex (Fig. S23). 85% of the amide signals for the [^15^N, ^13^C] L-CXCL8/D-3A11 complex were assigned. The chemical shift perturbation (CPS) profiles of the assigned amide signals of [^15^N, ^13^C] L-CXCL8 followed a similar pattern upon binding D-2A5 and D-3A11 peptides, with the exception for those of Arg68 (Fig. 3F). The binding of both peptides caused perturbations of Asp52, Val61, and the *C*-terminal α-helix which are characteristic for the CXCL8 dimer dissociation^51^. Positively charged residues involved in the GAG binding^52^ – Lys15, Lys20, Lys23, and Lys64 - were also significantly perturbed upon binding of peptides. In contrast, residues of the N- and 30s loops, which are responsible for the binding of the CXCR2 receptor^53^, remained unaffected. CXCL8 activity depends on an intricate equilibrium between dimerization, interaction with GAGs, and receptor binding^54^. To test whether selected D-peptides can disrupt this interplay, we tested them in migration and GAG-binding assays. The D-peptides did not inhibit CXCL8-induced migration of Jurkat cells at 10 μM concentration but effectively prevented binding of CXCL8 to GAGs (Fig. 4A-B).

**Figure 4.**
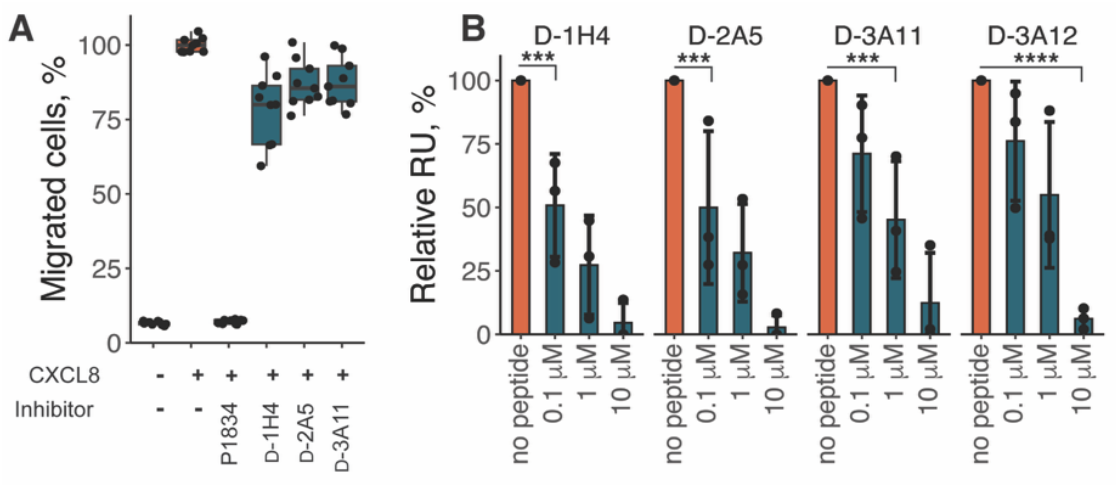
**A**. CXCL8-induced migration of Jurkat cells in the presence of 10 μM D-peptides. Values are normalized to migration induced by EC80 concentration of CXCL8, 1 μM of tick evasin P1834 is used as a positive control; all experiments are performed in triplicate in three biological replicates. B. CXCL8 binding to heparin in the presence of D-peptides. ^***^ - p ≤ 0.001 ^****^ - p ≤ 0.0001,

In summary, cyclic CLIPS™ L-peptides binding D-CXCL8 with sub-micromolar affinity were successfully selected using mirror-image phage display. We showed that the corresponding CLIPS™ D-peptides disrupt L-CXCL8 dimers and inhibit L-CXCL8 binding to GAGs, but not chemokine-mediated cell migration. Although blocking of chemokine-GAG interaction has been proposed as an alternative strategy to chemokine neutralization^55^, selected peptides require significant structural optimization to achieve binding affinities that bring therapeutic applications within reach. The authors face promising perspectives given recently published data on affinity-improvement of a PCSK9-inhibiting lead CLIPS™ peptide^56,57^ with very similar (i.e. submicromolar) affinity as observed here. Affinity improvement could be achieved by introducing structurally diverse D-amino acids (β1-, β2-, homo-, N-Me, and α-Me variants) currently available as building blocks for peptide synthesis, changes in the size of peptide’s binding loop (amino acids deletion and addition), as well as variations in the cyclisation approach.

## Acknowledgements

This work is supported by: MUMC+ Kootstra Fellowship and NWO-Rubicon 019.202EN.038 to S.S.D., NWO-ECHO 711.018.005 to I.D., KU Leuven C1 grant C14/23/143 to P.P., FWO-Vlaanderen PhD fellowship 1SHBW24N to F.B., Global PhD Partnerships - KU Leuven & Maastricht University GP-22-00193 to E.L.B., Investment Grant 91107006 to Dr. Gerry A.F. Nicolaes, Maastricht University.

## Methods

### Boc-SPPS general procedures

Boc-protected amino acids and resins were purchased from Bachem, Novabiochem, or Iris Biotech. Peptides were synthesised manually by iterative Boc-SPPS on Pam or MBHA resin. In short, each cycle consisted of a deprotection step of two 1-min. trifluoroacetic acid (TFA) treatments, 10 - 20 min treatment with a pre-activated amino acid followed by DMF washing. 1 eq. of the resin was swelled in DMF, and 4 eq. of Boc-protected amino acids activated by 4 eq. of HCTU (Peptides International, the USA) and 11 eq. of DIPEA were added to the resin containing the growing peptide chain. Coupling of Gln residues included an additional washing step by DCM to avoid heating and subsequent intramolecular pyrrolidone formation. Deprotection of Xan-protected residues was carried out in the presence of 5% triisopropylsilane (TIS) to prevent reaction of the xanthyl protective group with Trp residues. His was coupled in the Dnp-protected form. Eventually, the Dnp protection group was removed by thiophenol and benzyl mercaptan simultaneously with native chemical ligation (NCL). After completion of the peptide chain, peptides were deprotected and cleaved from the resin by anhydrous HF treatment for 1 h at 0 °C in the presence of 4% *p*-cresol as a scavenger. Crude peptides then were precipitated by ice-cold diethyl ether, dissolved in acetonitrile (ACN)/H_2_O (90:10 v/v) containing 0.1% TFA and lyophilised.

Preparative and semi-preparative RP-HPLC was performed using the Waters Deltaprep System consisting of a Waters Prep LC Controller and a Waters 2487 Dual wavelength Absorbance Detector equipped with a Vydac C18 22×250 mm (20 mL/min flow rate) or Vydac C18 10×250 mm (12 mL/min flow rate) HPLC column. Peptides **5** and **6** were purified using a Vydac C18 HPLC 4.6×150 mm column at 1 mL/min flow rate connected to a Varian Prostar system consisting of two Varian Prostar 215 delivery modules and a Varian Prostar 320 UV/Vis detector. Linear gradients of 0.1 % TFA in H_2_O (eluent A) and 0.1% TFA in ACN/H_2_O (90:10 v/v) (eluent B) were used for peptide and protein separation. Detection was carried out using absorbance at 214 nm.

LC-MS analysis was performed on a Waters UHPLC XEVO-G2QTOF system equipped with a C18 column. The mobile phase consisted of 0.1 % formic acid (FA) in H_2_O (eluent A) and 0.1% FA in ACN/H_2_O (90:10 v/v; eluent B). Molecular masses were calculated by deconvolution of mass spectra using MaxEnt 3.0 software (Waters, USA).

### General procedure for Native Chemical Ligation (NCL)

C-terminal and N-terminal fragments were mixed at concentrations of 10 mg/mL, each in 0.1 M Tris-HCl, 6 M Gdn-HCl, 2% (v/v) thiophenol, 2% (v/v) benzyl mercaptan, pH 8 and incubated at 37°C with intermittent mixing. Progression of ligation was followed by LC-MS. After completion of NCL, ligated material was purified by semi-preparative RP-HPLC, analysed by LC-MS, and lyophilised.

### D-CXCL8-biotin synthesis

#### Synthesis of ^34^D-Thz-D-Leu^49^-MPAL (1)

The middle part of the target protein was synthesised on 0.18 g of L-Leu-Pam resin (0.56 mmol/g, 100 μmol). Boc-D-Thioproline (Boc-D-Thz) was coupled as the last amino acid as an encrypted cysteine residue for further NCL. After cleavage from the resin, the peptide was lyophilised and purified by preparative RP-HPLC.

#### Synthesis of ^50^D -Cys-D-Ser^72^-KGG-biotin (2)

The C-terminal fragment was synthesised on 0.19 g of MBHA resin (0.53 mmol/g, 100 μmol). To allow coupling of biotin to the linker Boc-L-Lys(Fmoc)-OH was coupled as the first residue. Then Fmoc protection group was removed by four 3-min. treatment with 20% piperidine in DMF. After that, two Gly-residues were coupled to the lysine sidechain using HCTU-activated Fmoc-Gly-OH. Then HCTU-activated biotin was coupled to Gly-Gly linker. After completion of the biotin-linker, the rest of the peptide sequence was assembled. The resulting peptide was cleaved, lyophilised and purified by preparative RP-HPLC.

#### Synthesis of *N-terminus* ^*1*^*D-Ser-D-His*^*33*^*-MPAL (3)*

Similar to the synthesis of **1**, the synthesis of the N-terminal fragment was performed on 0.18 g of L-Leu-Pam resin (0.56 mmol/g, 100 μmol). Firstly, the resin was treated with HCTU-activated 3-tritylsulfanyl-propionic acid (Trt-MPA) for 30 min. After the removal of the trityl protecting group by 95% TFA, 2.5% TIS, 2.5% H_2_O, the rest of the peptide chain was assembled. After completion of the synthesis, the peptide was cleaved from the resin, precipitated with ice-cold diethyl ether, lyophilised and purified by preparative RP-HPLC.

### NCL of segments 1 and 2

^50^D-Cys-D-Ser^72^-KGG-biotin **2** (2.6 mg, 0.8 μmol) and ^34^D-Thz-D-Leu^49^-MPAL **1** (1.8 mg, 0.9 μmol) fragments were ligated according to the general procedure. After the ligation was complete (∼ 5h), the ligation mixture was diluted with 20 mL of 6 M Gdn-HCl, 0.1 M acetate buffer (pH 4) supplemented with 0.1 M *N*-methylhydroxylamine and mixed overnight at RT. The ligated fragment **4** was purified by semi-preparative RP-HPLC and lyophilised. Yield: 0.8 mg.

### NCL of segments 3 and 4

^1^D-Ser-D-His^33^-MPAL **3** (1.5 mg, 0.34 μmol) and ^34^D-Cys-D-Ser^72^-KGG-biotin **4** (0.8 mg, 0.16 μmol) were ligated according to the general procedure. After 2 h the ligation was complete, the full-length reduced D-CXCL8-KGG-biotin **5** was purified by semi-preparative RP-HPLC and lyophilised. Yield: 0.8 mg.

### Oxidative folding of reduced D-CXCL8-KGG-biotin 5

0.4 mg of lyophilised reduced D-CXCL8-KGG-biotin **5** protein was dissolved in 100 µL of 6 M Gdn-HCl, 0.1 M Tris buffer (pH 8) and then added dropwise to 0.9 mL of 1 M Gdn-HCl, 0.1 M Tris, 10 mM cysteine, 1 mM cystine buffer (pH 8). The progress of refolding was followed by LC-MS analysis. After completion of the conversion, folded D-CXCL8-KGG-biotin **6** was purified by semi-preparative RP-HPLC and lyophilised. Yield: 330 μg.

### Chemokines

Synthesis of the metabolically enriched [^13^C, ^15^N] L-CXCL8, L-CXCL8-biotin, L-CXCL1-biotin, and L-CCL5-biotin was described previously^16,50,58^. L-CXCL4-biotin was kindly provided by Dennis Suylen (Maastricht University). CXCL8 and CCL5 for migration and GAG-binding assays were purchased from Peprotech

### Phage-display selection

A phage display library comprising CLIPS™ peptides of the format AC_T2_XXXXXXXXXXC_T2_G (C_T2_ = cysteine cyclised via a bivalent T2-CLIPS™ scaffold, × = any amino acid excluding cysteine) as an N-terminal fusion with gIIIp was generated previously using the pADL^™^-100 phagemid vector (Antibody Design Laboratories, the USA) following the manufacturer protocol. Phage production and selection were performed as reported previously^59^ with a few modifications. In short, E. coli phage library glycerol stock was inoculated in a 2 L Erlenmeyer flask with 400 mL 2xYT, 100 µg/mL ampicillin, 2% glucose until OD_600_ reached 0.05. At an OD_600_ of 0.5 Hyperphage M13 KO7ΔpIII (Progen, Germany) was added according to manufacturer instructions. The culture was incubated at 37 °C for 30 min without shaking, then for 45 min at 250 rpm, and finally was centrifugated at 2000 g, 4 °C for 15 min. The cell pellet was resuspended in 400 mL 2xYT, 100 µg/mL ampicillin and 50 µg/mL kanamycin and was incubated overnight at 30 °C and 250 rpm. The overnight culture was centrifuged at 4000 g and 4 °C for 30 min and the supernatant was harvested. Phages were precipitated by the addition of 80 mL of ice-cold 20% polyethylene glycol (PEG-8000) in 2.5 M NaCl and incubation on ice for 4 h. Next, the phage-suspension was centrifugated at 10,000 g and 4 °C for 30 min, the supernatant was discarded, and the phage pellet was resuspended in 1 mL of TBS buffer (20 mM Tris, 150 mM NaCl, pH 7.4). The phage solution was centrifuged at 5400 g for 15 min at 4° C and sterile filtered (cut-off: 0.45 µm) to remove any remaining bacterial cells.

For cross-linking of peptides displayed on phages with 1,3-bis(bromomethyl)benzene (bivalent T2-scaffold), phages were first reduced by addition of 200 µL of ammonium bicarbonate (ABC) buffer (440 mM, pH 8.5), 7.5 µL of TCEP (100 mM) and 7.5 µL of EDTA (500 mM) to 500 µL of the phage-solution and incubation for 1 h at 37 °C. After that, 10 µL of 10% acetic acid was added and the phage-solution buffer-exchanged with 2 mL of milliQ-H_2_O and an MWCO 40k Zeba™ spin column (ThermoFisher Scientific). The reduced phage solution was added to 90 µL of the bivalent T2-scaffold (500 µM in ACN), mixed and then 90 µL of ABC-buffer (440 mM, pH 8.5) was added to cyclise the peptides with the bivalent T2-scaffold for 1 h at 30 °C. After that, 7.5 µL of cysteine (50 mM) was added and the solution was incubated for an additional 15 min to quench the excess T2-scaffold. Finally, the phage solution was buffer exchanged with 2 mL of PBS using MWCO 40k Zeba™ spin column and was further used for the phage selection of CLIPS™-cyclised peptides.

For phage-display selection, 2.5 µg of D-CXCL8-biotin and 2.5 µg of BSA-biotin (Abcam, the UK) were captured on 100 µL Dynabeads® M-280 Streptavidin magnetic beads (ThermoFisher Scientific) in 1 mL PBS for 30 minutes and non-immobilised protein was removed by washing three times with PBS, 0.05% Tween-20 (PBST). The beads were then blocked under inversion with 1 mL of 4% BSA in PBS for 1 h. After that, the blocked beads were again washed three times before use in the selection procedure.

100 µL of the magnetic beads coated with BSA-biotin were resuspended in 100 µL of BSA (10%) in PBS, then 900 µL of the cross-linked phage-solution in PBS was added, mixed and incubated for 1 h under inversion to remove sticky or bead-binding peptides from the phage library. Then, the beads were removed, and the supernatant of the phage solution was added to the beads coated with the target protein D-CXCL8-biotin. The bead-phage suspension was inverted for 1 h at room temperature. After that, the supernatant was discarded and the beads were washed 2 and 5 times with 1 mL of PBS-Tween (PBST) for rounds 1 and 5, respectively, and one final time with 1 mL of PBS. Finally, the phages were eluted from the beads by addition of 0.8 mL triethylamine (100 mM) and mixing for 5 min. Then, the supernatant containing eluted phages was recovered and quenched by the addition of 0.5 mL of 1M Tris-HCl, pH 7.4 and then added to 25 mL of an exponential culture of TG1 cells in 2xYT with 1% glucose. The culture was incubated in a shaker at 37 °C for 30 min. without and 30 min. with shaking at 50 rpm. Finally, 25 µL of 100 mg/mL ampicillin was added and the culture was incubated overnight at 30 °C and 250 rpm. On the next day, a glycerol stock was prepared from the overnight culture and fresh phages were produced for the next round of phage display starting, as described above, by inoculating media with cells at an OD_600_ of 0.05. In total three rounds of phage display selection were performed to enrich the phage pool in binders for D-CXCL8.

### NGS data and bioinformatic analysis

After the 3^rd^ round of selection, the TG1 culture was harvested, and the DNA of selected phages was isolated with the GeneJET Plasmid Miniprep Kit (ThermoFisher Scientific). The purified plasmid pool was sequenced by Illumina NovaSeq6000 with a 150 bp reading frame (>5 million paired-end reads per sample) using the commercial service provided by Genomescan B.V, the Netherlands. Data were processed by Genomescan B.V. using custom R-scripts to filter and retain reads that match the correct DNA reading frame and format of the phage library. Retained DNA sequences were then translated into peptide sequences, and for each unique peptide sequence, the abundance in the NGS population was calculated. The 500 most abundant sequences in the NGS population were then further analysed as described in bioinformatic analysis.

Data manipulation and visualisation were performed using R 4.1.1 with specified packages. The variable region of the 500 most abundant sequences from the NGS data were aligned by *Biostrings_2*.*62*.*0* package using the “Muscle” method with the gap penalty of 999. The chemical space of obtained peptides was visualised by calculating of the pair-wise dissimilarity matrix for aligned sequences and subsequent dimensionality reduction by metric multi-dimensional scaling using *bios2mds_1*.*2*.*3*. K-means clustering into 4 groups was performed using *factoextra_1*.*0*.*7* and visualised by *ggplot2_3*.*3*.*6*. Data for pLogo plots for each cluster were generated using the web server^40^ with the naive phage library before selections as a reference data set. Subsequently, plots were created with *ggseqlogo_0*.*1* using only overrepresented amino acids. Peptide properties were calculated using *Peptides_2*.*4*.*4*. Sequence identity was calculated using pair-wise alignment as 100×(identical positions)/(aligned positions + internal gap positions) by *Biostrings_2*.*62*.*0*.

### CLIPS™ peptide synthesis

Synthesis of CLIPS™-cyclised peptides was performed as described previously^60^. The starting linear peptides were synthesised on fully automated peptide synthesisers (Multisyntech, Syro, 2 μmol scale for crude libraries) or Gyros Protein Technologies (Symphony, 100-200 μmol scale, for bulk production) using Fmoc-based solid-phase peptide synthesis on TentaGel^®^ Ram resin using standard protocols. Coupling of L- and D-cysteines was performed using 2,4,6-trimethylpyridine (TMP) as a base to maximally suppress racemisation. Peptides (100-200 μmol) were purified by preparative HPLC on a Reprosil-Pur 120 C18-AQ 150×20 mm column (Dr. Maisch GmbH, Germany) using an ACN/H_2_O gradient (5-65%) containing 0.05% TFA followed by lyophilisation on an Alpha 2-4 LD plus (Martin Christ Gefriertrocknungsanlagen GmbH, Germany). The 2 μmol peptide libraries were used without further purification. Lyophilisation was performed using a GeneVac HT-4X centrifugal vacuum evaporator. The peptides were then dissolved in DMF (0.5 mL), T2-scaffold dissolved in DMF (4.1 mM, 0.5 mL) was added and the solution was homogenised, followed by the addition of ABC (150 mM, pH 8.0, 0.5 mL) immediately followed by homogenising the resulting solution. After 1 h at room temperature, the reaction mixtures were quenched with 0.5% ethane thiol (in 1:1 DMF/H_2_O, 0.1 mL/peptide). Finally, the CLIPS™-cyclised peptide libraries were then lyophilised at least 3x using a Genevac HT-4X evaporation system.

Synthesis of purified CLIPS™-peptides was carried out with a 0.5 mM solution of the linear peptides in ACN/H_2_O (1:3, v/v) to which was added 1.1 equiv. of 1,3-bis(bromomethyl)benzene (bivalent T2-scaffold) dissolved in ACN, whereafter the solution was homogenised. Then, 44 equiv. of ABC (0.2 M) were added, and the solution was homogenised and reacted for 60 min. After completion (monitoring by UPLC), the reaction was quenched with 10% TFA/H_2_O to pH < 4 and directly loaded onto a preparative RP-HPLC, or first lyophilised and subsequently purified by preparative RP-HPLC.

### Surface Plasmon Resonance (SPR) biosensor assay

Library screenings, titrations and selectivity experiments were carried out using Biacore T200 (Cytiva) with SAHC200M sensor chips (XanTec bioanalytics GmbH, Germany). For the library screening and Ala-scanning experiment, 3 kRU of biotinylated L- and D-CXCL8 have been immobilised on the sensor surface. To eliminate peptides with high non-specific binding to the matrix of the chip, all peptides were injected for 60 s over the reference non-modified surface at the nominal concentration of 10 µM in phosphate buffer pH 7.4 supplemented with 0.05% Tween-20, 30 µL/min. Peptides with non-specific binding higher than 200 RU were eliminated from the screening. The remaining peptides were injected against D- and L-CXCL8 channels for 60 s, after each injection 30 s pulse 0.5 M formic acid was applied as a regeneration step.

For titration and selectivity experiments SAHC200M chips with lower immobilisation levels were used: ∼1.5 kRU of D-and L-CXCL8 or 1.5 kRu of L-CXCL1, L-CXCL4, and L-CCL5, respectively. 1 mM DMSO stock solutions of peptides were diluted by the running buffer to final concentrations of 2 µM – 15 nM. 1 % DMSO was added to the running buffer to match DMSO content with the sample buffer. Samples were injected for 180 s at 30 µL/min followed by 240 s of dissociation time. 30 s injection of 0.5 M formic acid was used as a regeneration step. Each sample injection was followed by an injection of the running buffer as a chip surface conditioning step. Sensograms were analysed by BioEvaluation software. K_D_ values were calculated using in-built affinity analysis with the 1:1 binding model as an average of three independent measurements.

Competition for binding to heparin was evaluated using a Biacore 8K (Cytiva) and a CM4 Series S sensor chip (Cytiva). Immobilization of biotinylated heparin on a surface was carried out at 10µL/min as follows. The chip was equilibrated in HBS-EP+ Running Buffer (10 mM HEPES, 150 mM NaCl, 3 mM ethylenediaminetetraacetic acid (EDTA), 0.05% P20 surfactant, pH 7.4), then regenerated 3 times in 1 M NaCl, 50 mM NaOH for 60 s, activated by a 1:1 mixture of 0.05 M N-hydroxysuccinimide (NHS) and 0.2 M (N-(3-dimethylaminopropyl)-N’-ethylcarbodiimide hydrochloride) (EDC) for 900 s, before finally washing the fluidics system with ethanolamine. Then, a 20 µg/mL solution of NeutrAvidin (#31000; Thermo Fisher Scientific) in 10 mM sodium acetate (pH 5.5) was injected for 30s until an immobilization level reached ∼200 absolute RU. Unreacted carboxyl groups were quenched with 100% ethanolamine for 900 s. Subsequently, ∼ 4-7 relative RU of biotinylated low molecular weight heparin (4.5 kDa) were immobilized in an active chip lane by injection of either a 445 nM or a 200 nM solution in HBS-EP+ Running Buffer for 60 s.

GAG-binding screening was performed by injecting a 100 nM solution of CCL5 (#300-06; PeproTech) or CXCL8 (#200-08; Peprotech) in increasing concentration of peptides in HBS-EP+ Running Buffer + at 30µL/min for 200 s. The signal was double referenced and normalized to the signal obtained by injection of 100nM chemokine in the absence of peptides. Extensive washing was performed between each cycle with 1M NaCl (contact time 30s, flow rate 30µL/min), and three regeneration steps with 1M NaCl + 20mM NaOH were performed at the beginning of each separate experiment (contact time 30s, flow rate 30µL/min). Data were analyzed with Cytiva Biacore™ Insight Evaluation Software.

### NMR spectroscopy

NMR spectra were recorded using Bruker Avance III HD 700 MHz spectrometer, equipped with a cryogenically cooled TCI probe as described previously^16,50^.

Spectra processing was performed by Bruker Topspin 3.2 and NMRFAM-SPARKY 3.114 software. Secondary structure prediction for peptides was performed using CSI 3.0 server^47^. Dihedral angles were predicted using TALOS^61^ and inverted for the structure calculation. The structure calculations were performed using XPLOR-NIH^49^ with the eefx force field^48^. The Ramachandran plots were plotted using an inverted Top8000 reference data from the Richardson lab (https://github.com/rlabduke/reference_data/).

### Jurkat:CXCR1 cell migration assay

Jurkat E6.1 cells (ATCC TIB-152) were transfected by electroporation with PvuI-linearised plasmid D1398 encoding human CXCR1. Transfected clones were selected, expanded and maintained in RPMI-1640 (R0883, Sigma), 10% FBS (F9665, Sigma), 5 mM L-Glutamine (G7513, Sigma) supplemented with 5 μg/mL blasticidin (203350, Sigma). The Jurkat:CXCR1 cells were used for generating dose-response curves with ELR+ chemokines. Peptides were tested against the EC80 concentration of the CXCL8 (0.81 nM). The migration assays were performed as described previously^18^. In short, 3×10^5^ ells/well were added to the top chamber of a 3-μm 96-well Transwell insert (3385, Corning) in 50 μL of cell migration media (RPMI-1640 (R0883, Sigma), 0.5% FBS (F9665, Sigma), 4 mM L-Glutamine (G7513, Sigma), 0.05% DMSO (D4540, Sigma)). The bottom chamber contained 150 μL of migration media with CXCL8 pre-incubated with D-peptide at 37°C for 30 minutes. Cells were migrated at 37°C in 5% CO_2_ for 2h. The migration plate was shaken at 850 RPM for 10 min, and 150 μL of media from the bottom chamber of the migration plate was transferred to a round bottom 96-well plate (353910, Falcon) containing 50 μL of migration media. Cell counts were determined using an Attune NxT Acoustic Focusing Cytometer with CytKick MAX AutoSampler (ThermoFisher).

### CD spectroscopy

CD spectra of 0.1 - 0.2 mg/ml L- and D-peptides were recorded with Chirascan V100 (Applied Photophysics, the UK) at 25 °C in ACN/H_2_O (45:55, v/v) containing 0.1% TFA, pathlength 1 mm, bandwidth 1 nm, step size 1 nm. Final spectra were averaged from three measurements after blank buffer subtraction using Chirascan v.4.7.0.194 software. Secondary structure determination was performed using BeStSel server^45^.

