## Supplemental Information for "Chemokine-binding all-D-CLIPS™ peptides identified using mirror-image phage display"

**Table S1. Secondary chemical shifts and CSI 3.0 algorithm output for D-2A5 and D-3A11.**

B-prob, C-prob, and H-prob stand for the probability of β-sheet, coil, and α-helix, respectively.

**D-2A5**

| Seq | Cα | Cβ | H | Hα | B-prob | C-prob | H-prob |
|---|---|---|---|---|---|---|---|
| A1 | 51.61 | 20.04 | 8.04 | 3.883 | 0.14 | 0.86 | 0.00 |
| C2 | | 35.22 | 8.686 | 5.054 | 0.86 | 0.13 | 0.01 |
| W3 | 56.15 | 31.65 | 8.338 | 4.961 | 0.93 | 0.06 | 0.00 |
| V4 | 60.75 | 34.20 | 8.58 | 4.807 | 0.98 | 0.02 | 0.00 |
| I5 | 59.58 | 40.50 | 8.425 | 4.544 | 0.92 | 0.08 | 0.00 |
| S6 | | 64.65 | 8.429 | 5.019 | 0.77 | 0.23 | 0.00 |
| Y7 | 58.62 | 39.73 | 8.742 | 4.42 | 0.37 | 0.51 | 0.12 |
| D8 | 53.90 | | 8.762 | 4.128 | 0.18 | 0.82 | 0.00 |
| G9 | 45.88 | | 7.838 | | 0.08 | 0.85 | 0.07 |
| Y10 | 57.34 | 39.88 | 7.692 | 4.556 | 0.34 | 0.65 | 0.01 |
| E11 | 55.11 | 30.90 | 8.51 | 4.81 | 0.86 | 0.14 | 0.00 |
| Y12 | 57.12 | 40.86 | 8.588 | 4.692 | 0.83 | 0.17 | 0.00 |
| C13 | | 35.08 | 8.714 | 4.914 | 0.80 | 0.17 | 0.03 |
| G14 | 44.63 | | 8.209 | | 0.14 | 0.86 | 0.01 |

**D-3A11**

| Seq | Cα | Cβ | H | Hα | B-prob | C-prob | H-prob |
|---|---|---|---|---|---|---|---|
| A1 | 51.34 | 20.82 | | 4.066 | 0.37 | 0.63 | 0.00 |
| C2 | 55.30 | 36.02 | 8.56 | 4.692 | 0.79 | 0.19 | 0.02 |
| F3 | 56.81 | 42.00 | 8.66 | 5.092 | 0.94 | 0.06 | 0.00 |
| L4 | 54.99 | 45.53 | 8.536 | 4.491 | 0.85 | 0.15 | 0.00 |
| A5 | 51.07 | 21.14 | 8.355 | 4.813 | 0.92 | 0.08 | 0.00 |
| M6 | 55.05 | 35.28 | 8.601 | 4.571 | 0.70 | 0.29 | 0.01 |
| D7 | 53.41 | 39.23 | 8.97 | 4.539 | 0.09 | 0.91 | 0.00 |
| G8 | 45.15 | | 8.495 | | 0.12 | 0.86 | 0.02 |
| V9 | 61.27 | 34.69 | 7.784 | 4.147 | 0.69 | 0.31 | 0.00 |
| E10 | 54.88 | 30.58 | 8.242 | 4.622 | 0.63 | 0.37 | 0.00 |
| Y11 | 56.89 | 41.31 | 8.71 | 4.681 | 0.88 | 0.12 | 0.00 |
| R12 | 55.00 | 32.72 | 8.5 | 4.515 | 0.71 | 0.29 | 0.00 |
| C13 | 55.93 | 35.60 | 8.556 | 4.676 | 0.77 | 0.20 | 0.02 |
| G14 | 44.59 | | 8.568 | | 0.15 | 0.84 | 0.01 |

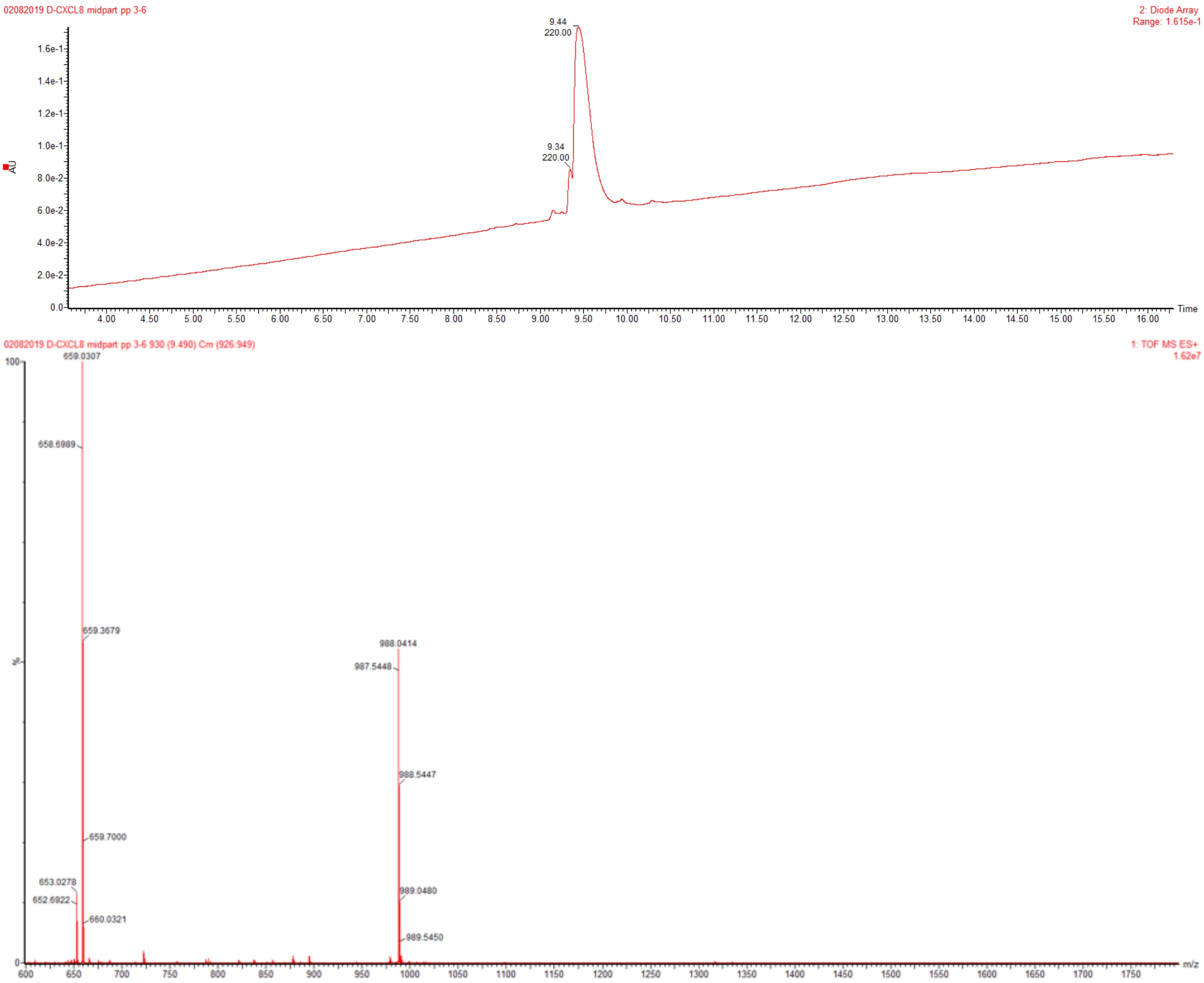

**Figure S1. LC-MS analysis of $^{34}$D-Thz-D-Leu$^{49}$-MPAL 1.**
The HPLC trace (top) and ESI mass spectrum (bottom) of 1. Observed deconvoluted $[M+H]^+$ MW = 1974.1 Da, calculated $[M+H]^+$ MW = 1973.9 Da.

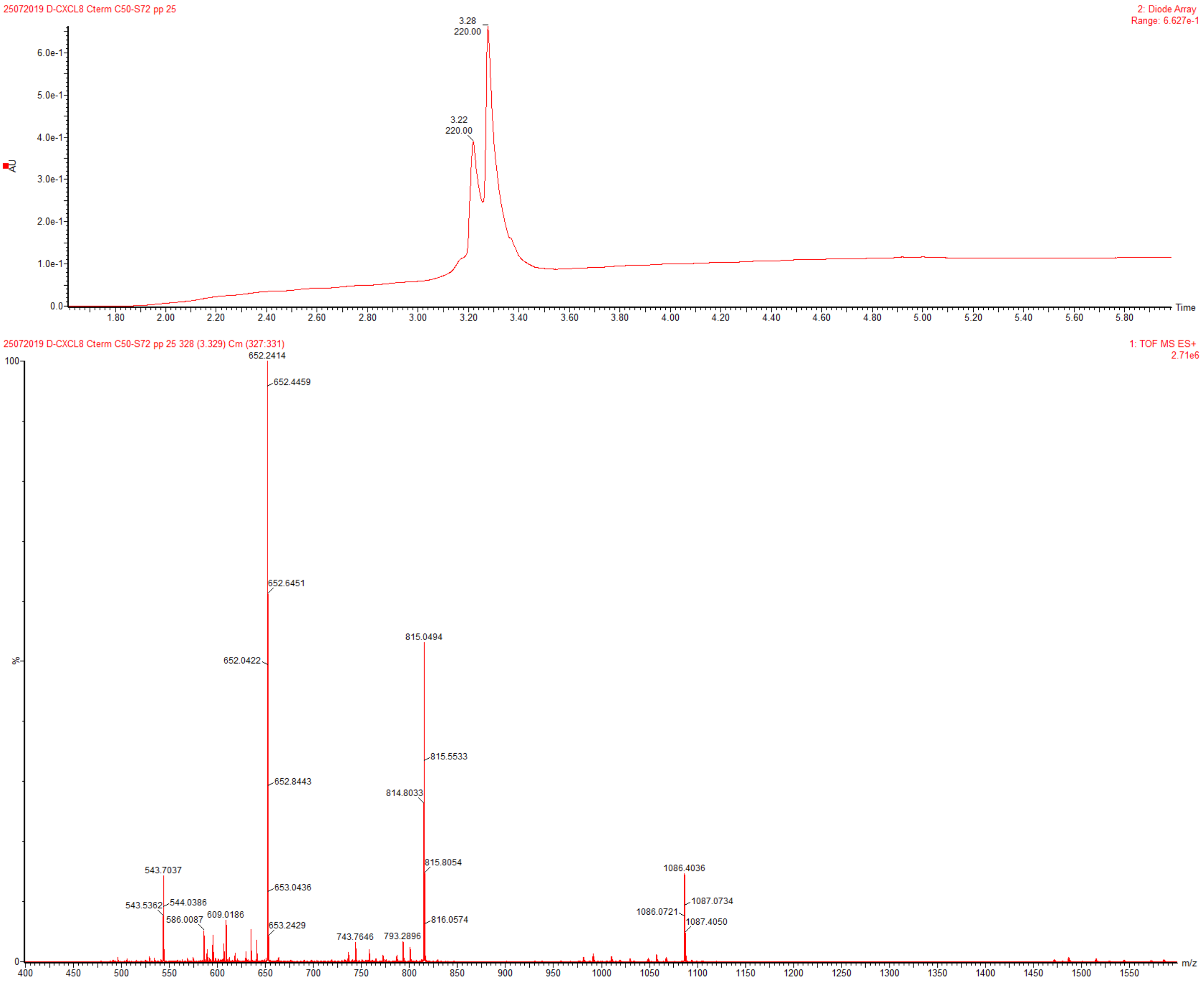

**Figure S2. LC-MS analysis of $^{50}$D-Cys-D-Ser$^{72}$-KGG-biotin 2.**

The HPLC trace (top) and ESI mass spectrum (bottom) of 2. Observed deconvoluted $[M+H]^+$ MW = - 3256.2 Da, calculated $[M+H]^+$ MW = 3255.9 Da. The minor peak is a Glu deletion.

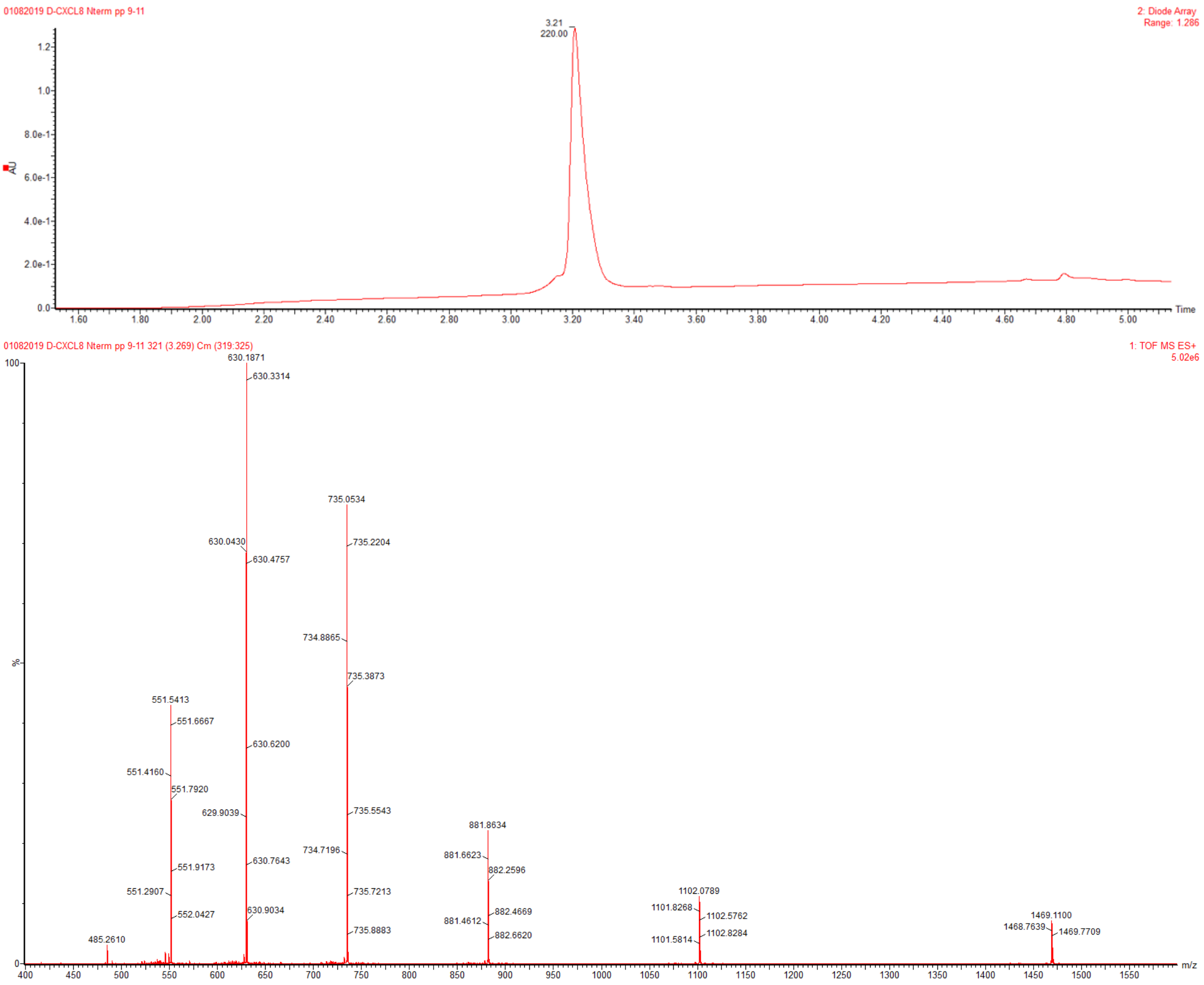

**Figure S3. LC-MS analysis of [1]D-Ser-D-His[33]-MPAL 3.**

The HPLC trace (top) and ESI mass spectrum (bottom) of 3. Observed deconvoluted $[M+H]^+$ MW = 4403.3 Da, calculated $[M+H]^+$ MW = 4403.2 Da.

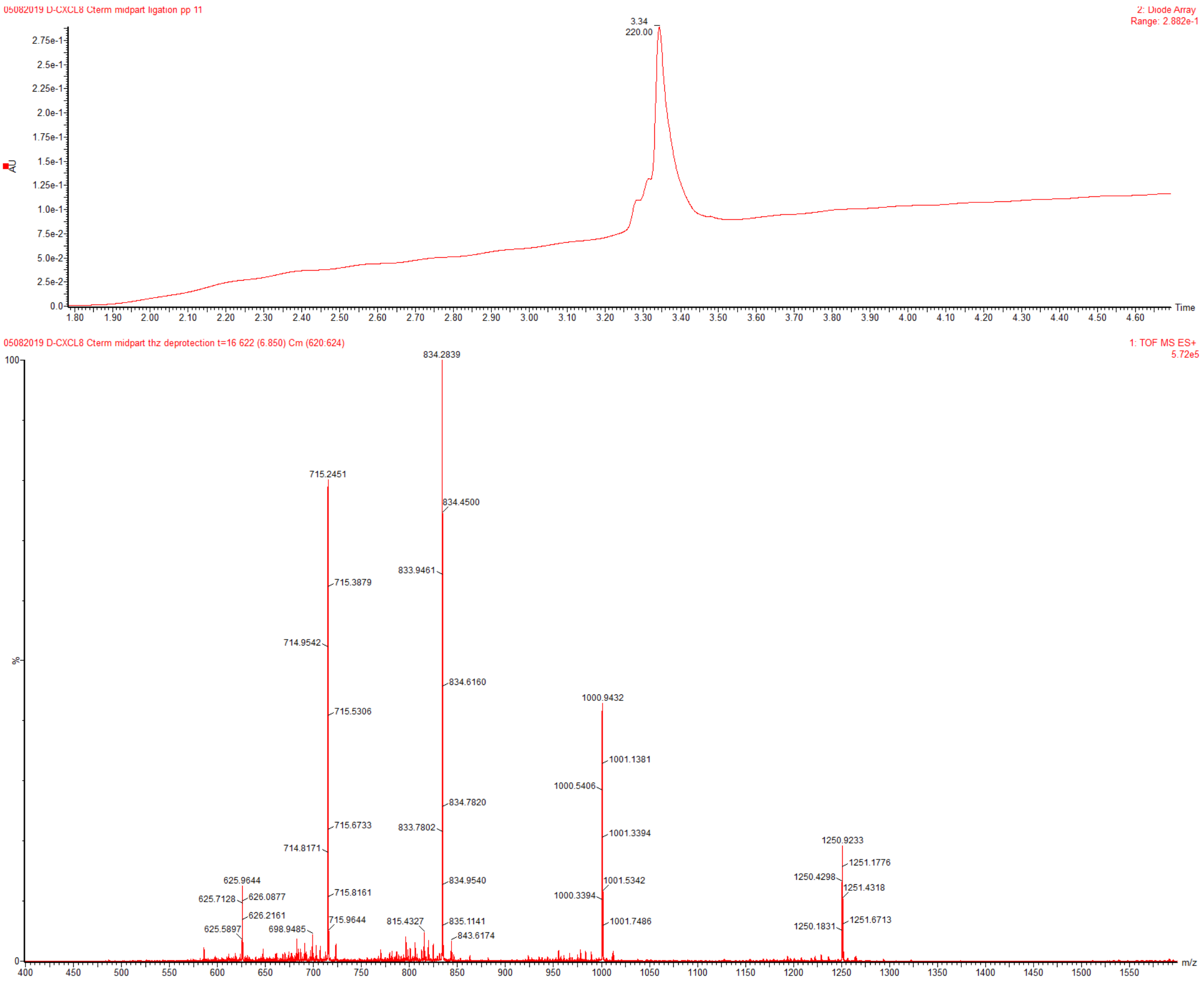

**Figure S4. LC-MS analysis of [34]D-Cys-D-Ser[72]-KGG-biotin 4.**

The HPLC trace (top) and ESI mass spectrum (bottom) of 4. Observed deconvoluted $[M+H]^+$ MW = 4997.6 Da, calculated $[M+H]^+$ MW = 4997.7 Da.

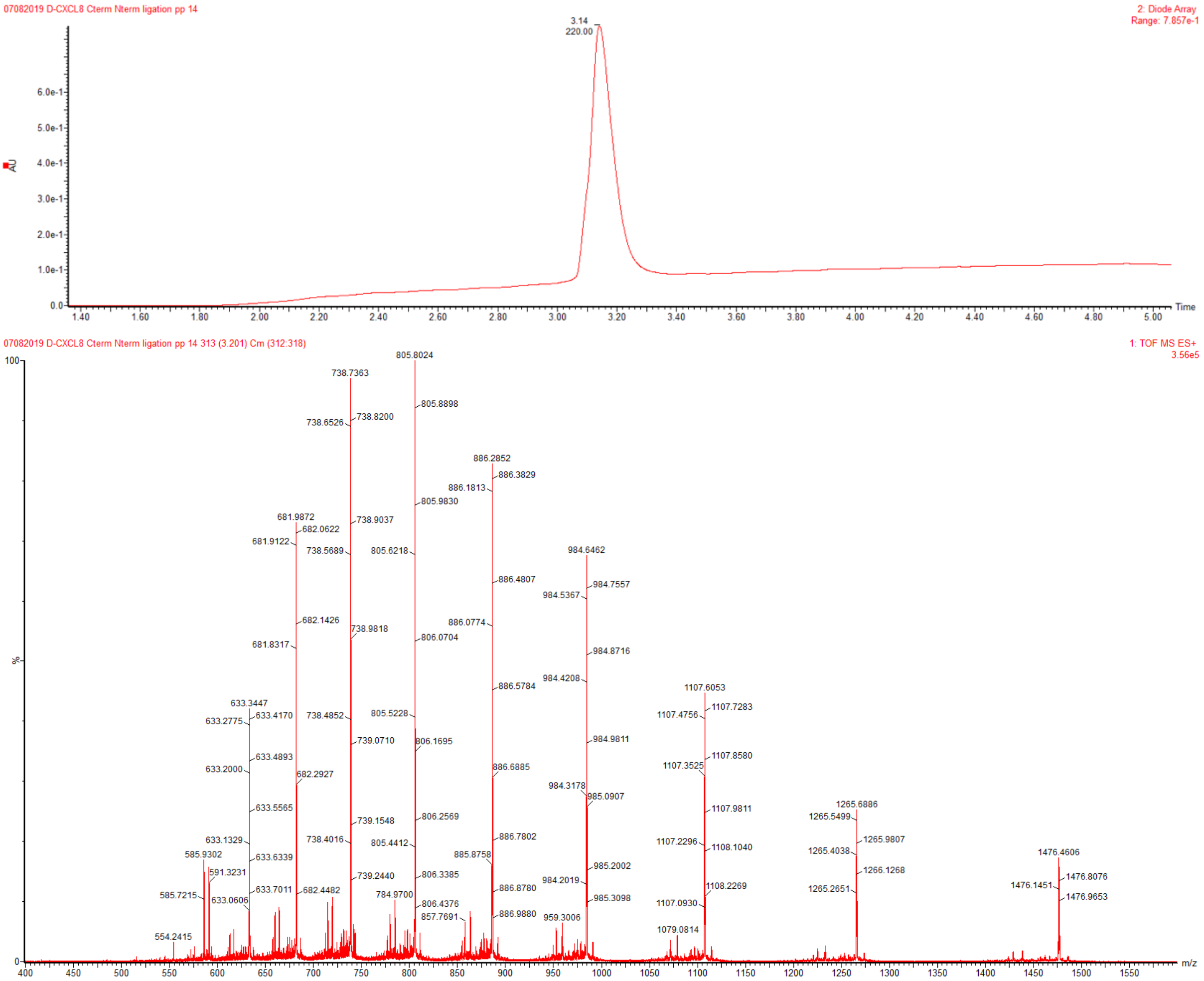

**Figure S5. LC-MS analysis of reduced D-CXCL8-KGG-biotin 5.**
The HPLC trace (top) and ESI mass spectrum (bottom) of 3. Observed deconvoluted $[M+H]^+$ MW = 8848.8 Da, calculated $[M+H]^+$ MW = 8848.8 Da.

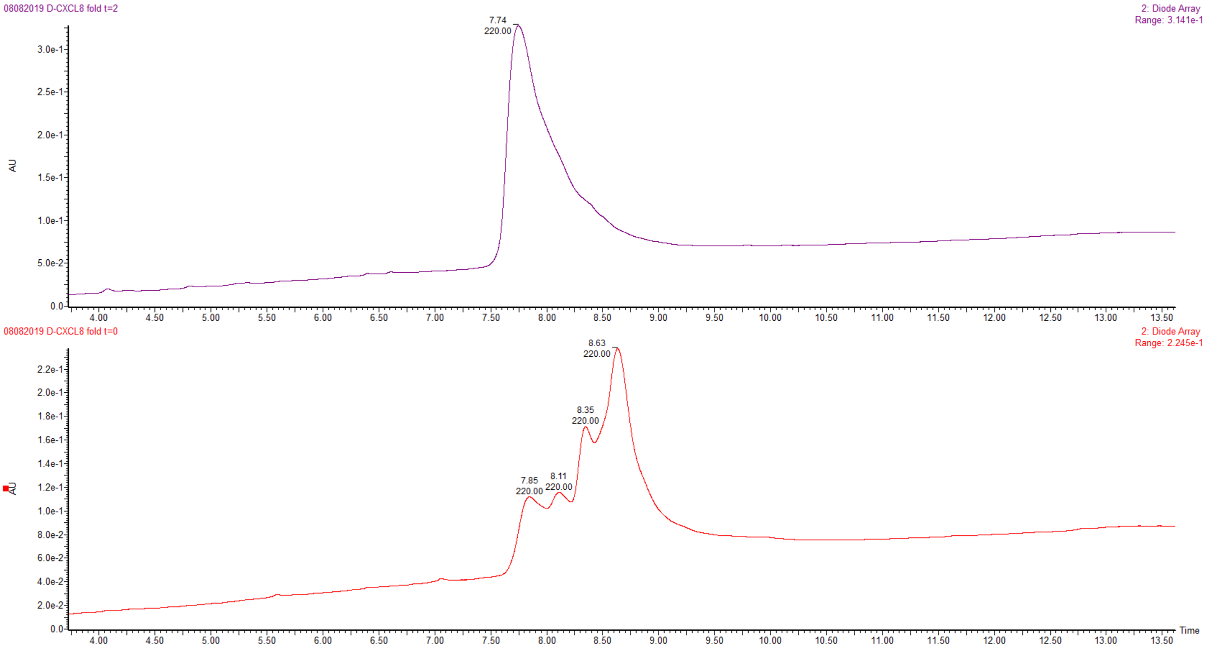

**Figure S6. D-CXCL8-biotin refolding**
HPLC traces of D-CXCL8-biotin under oxidative refolding conditions at t = 0 (bottom) and after 2 hours (top).

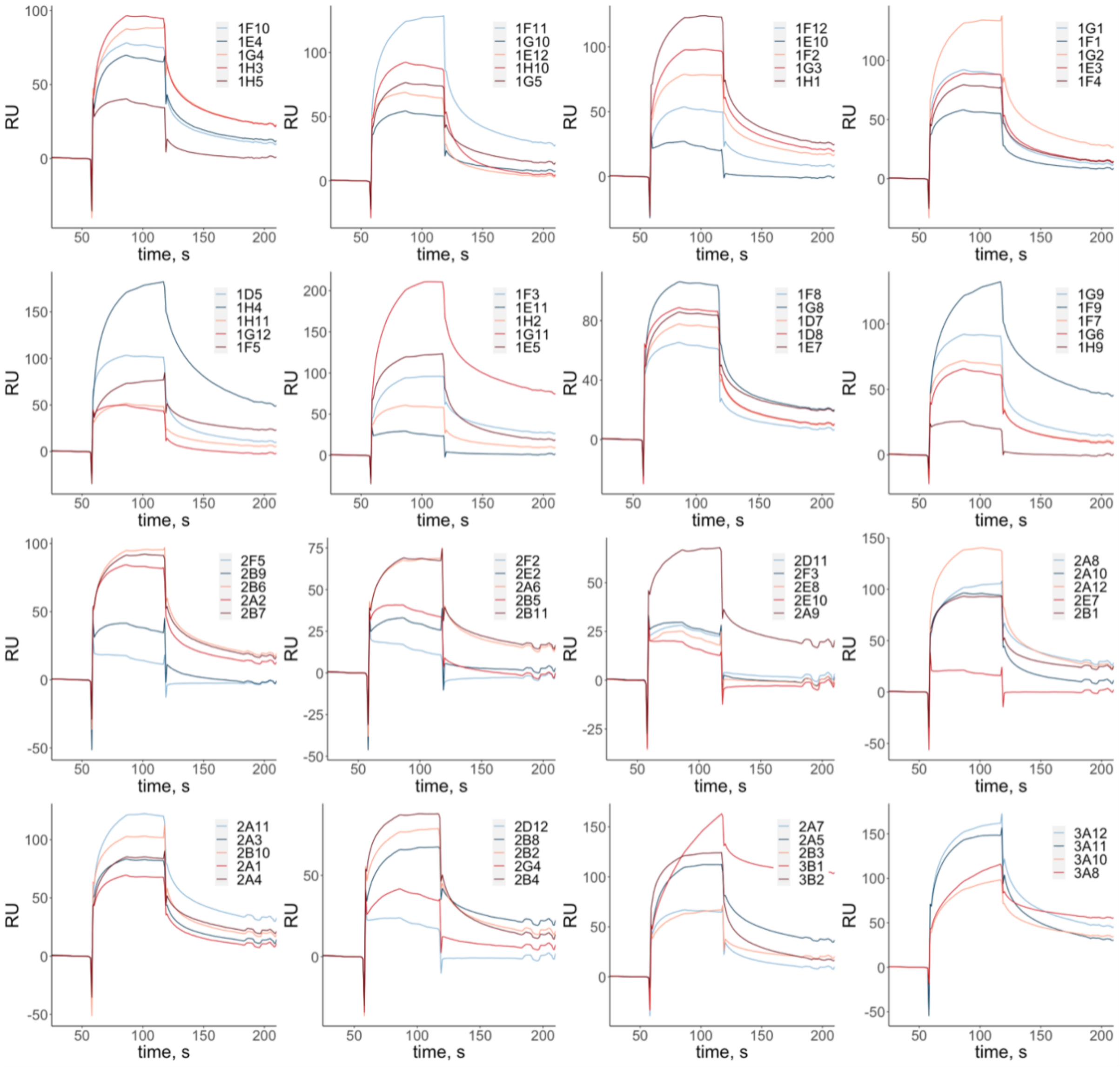

**Figure S7. Crude CLIPS$^{TM}$ L-peptide interaction with D-CXCL8-KGG-biotin (6).** SPR biosensor analysis of interaction of 10 μM crude L-CLIPS$^{TM}$ peptides with D-CXCL8-KGG-biotin (**6**).

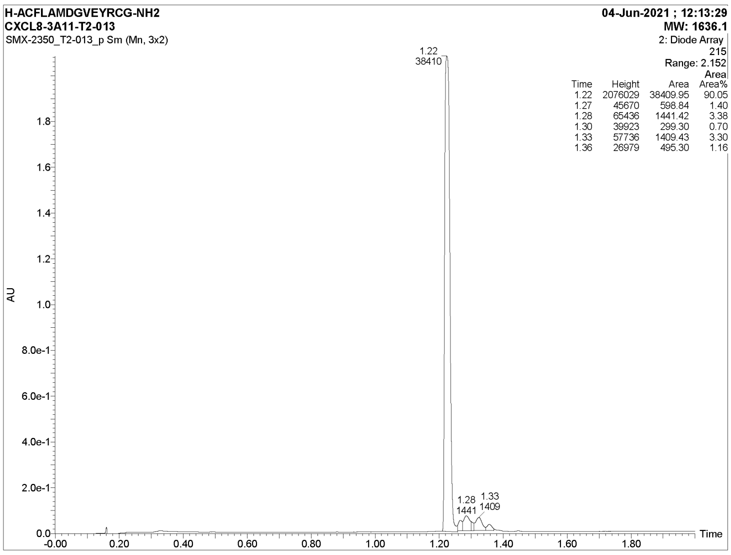

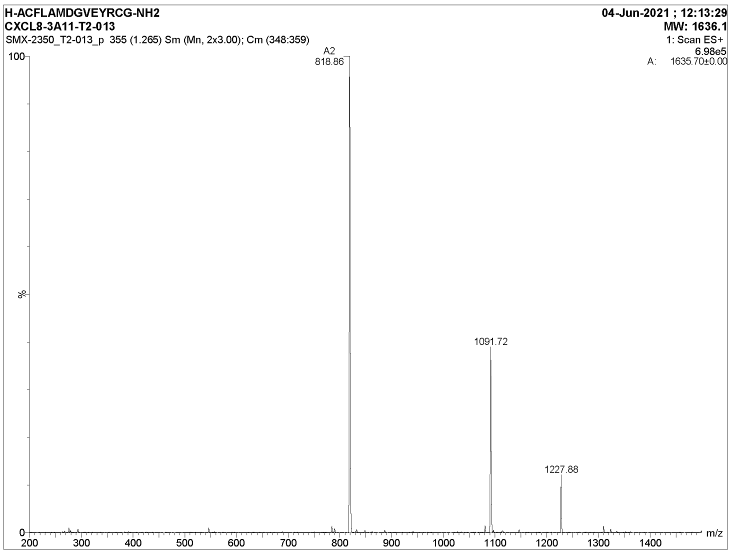

**Figure S8. LC-MS analysis of L-CLIPS$^{TM}$ peptide 3A11**

The HPLC trace (top) and ESI mass spectrum (bottom) of L-CLIPS$^{TM}$ peptide 3A11. Observed *m/z* signals: $[M+2H]^{2+}$ - 818.9, $[2M+3H]^{3+}$ - 1091.7, $[3M+4H]^{4+}$ - 1227.9; calculated *m/z* signals: $[M+2H]^{2+}$ - 819.1, $[2M+3H]^{3+}$ - 1091.7, $[3M+4H]^{4+}$ - 1228.1.

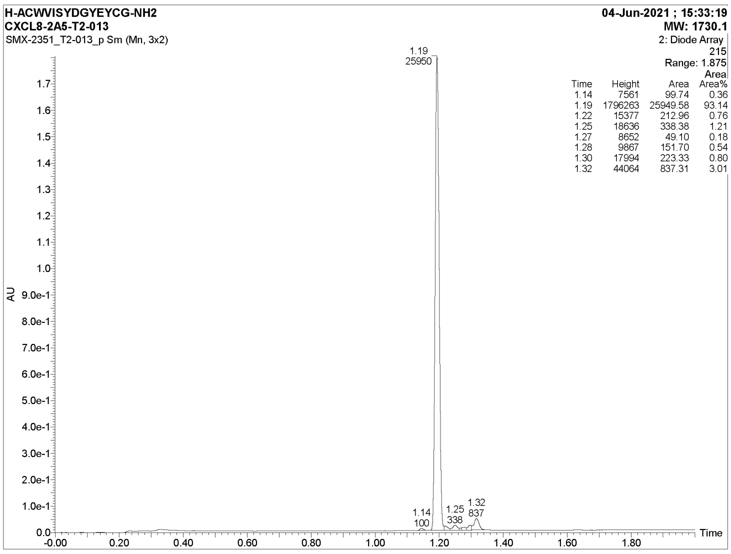

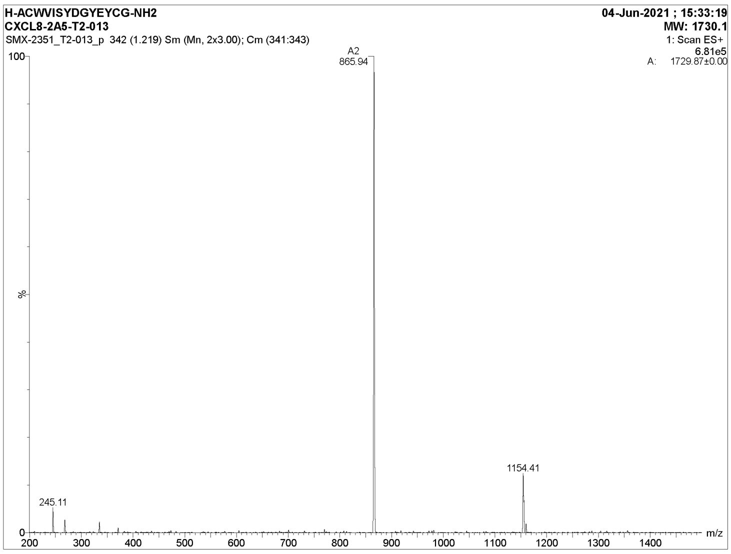

**Figure S9. LC-MS analysis of L-CLIPS™ peptide 2A5.**
The HPLC trace (top) and ESI mass spectrum (bottom) of L-CLIPS™ peptide 2A5. Observed *m/z* signals: $[M+2H]^{2+}$ - 865.9, $[2M+3H]^{3+}$ - 1154.4, calculated *m/z* signals: $[M+2H]^{2+}$ - 866.1, $[2M+3H]^{3+}$ - 1154.4.

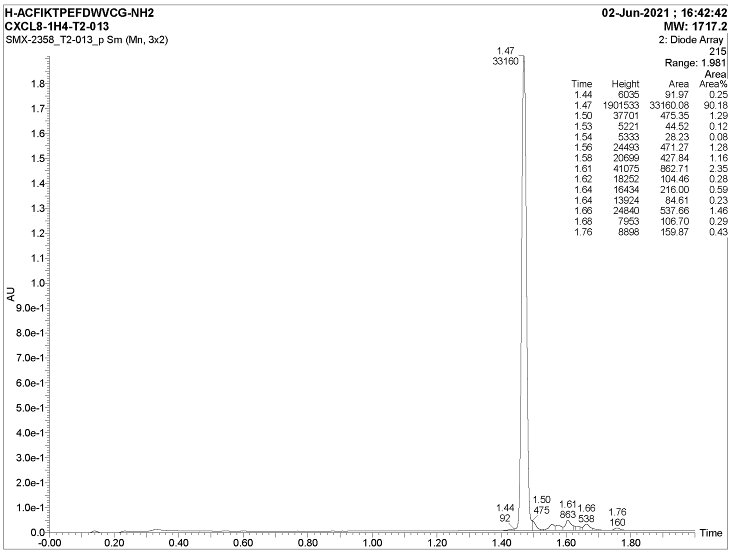

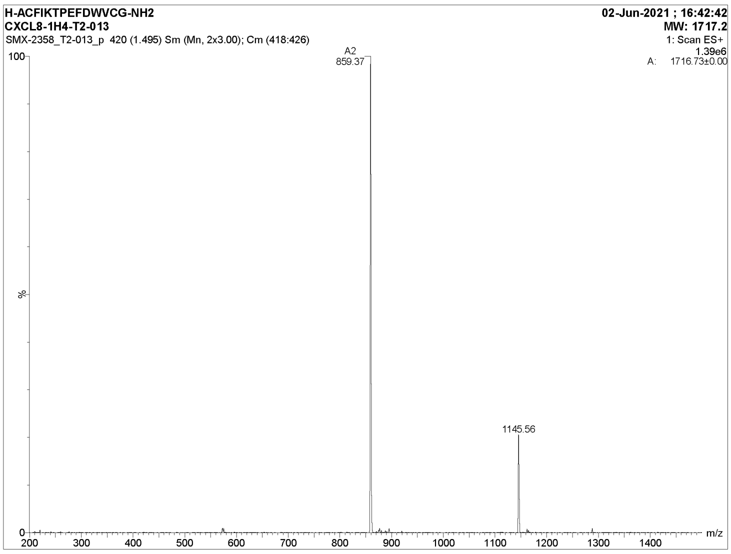

**Figure S10. Synthesis of L-CLIPS™ peptide 1H4.**

The HPLC trace (top) and ESI mass spectrum (bottom) of L-CLIPS™ peptide 1H4. Observed *m/z* signals: $[M+2H]^{2+}$ - 859.4, $[2M+3H]^{3+}$ - 1145.6, calculated *m/z* signals: $[M+2H]^{2+}$ - 859.6, $[2M+3H]^{3+}$ - 1145.8.

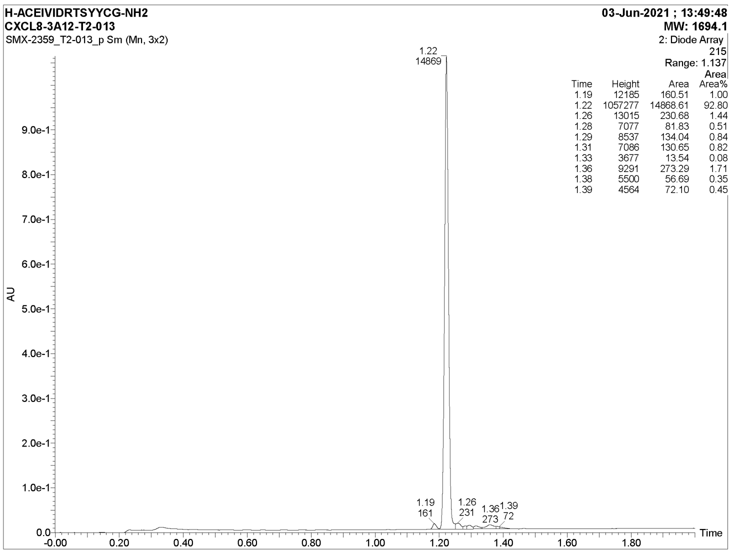

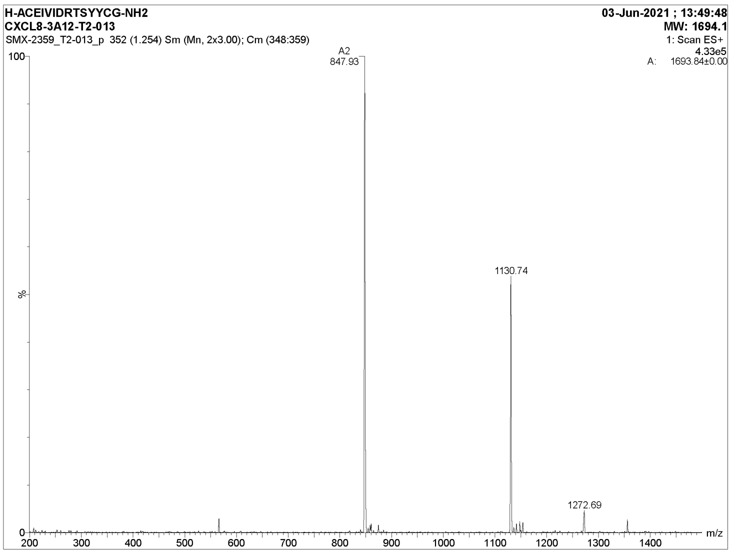

**Figure S11. LC-MS analysis of L-CLIPS™ peptide 3A12.**
The HPLC trace (top) and ESI mass spectrum (bottom) of L-CLIPS™ peptide 3A12. Observed *m/z* signals: $[M+2H]^{2+}$ - 847.9, $[2M+3H]^{3+}$ - 1130.7, $[3M+4H]^{4+}$ - 1272.7; calculated *m/z* signals: $[M+2H]^{2+}$ - 848.1, $[2M+3H]^{3+}$ - 1130.4, $[3M+4H]^{4+}$ - 1271.6.

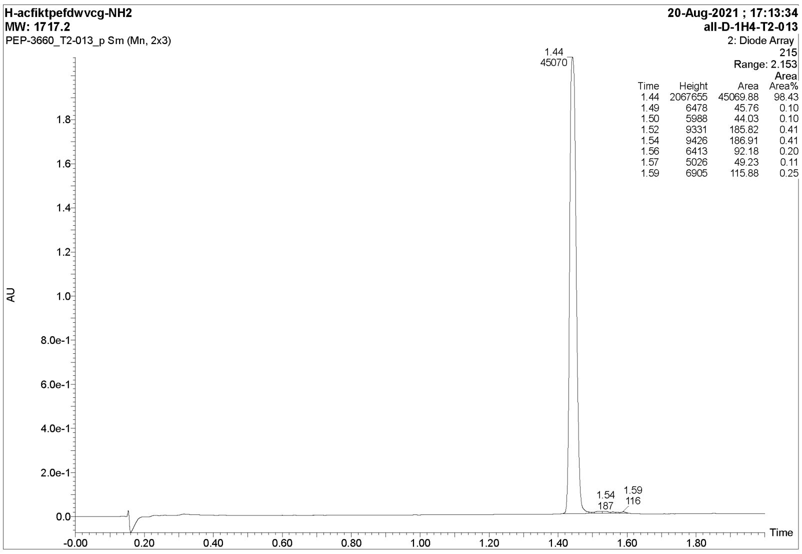

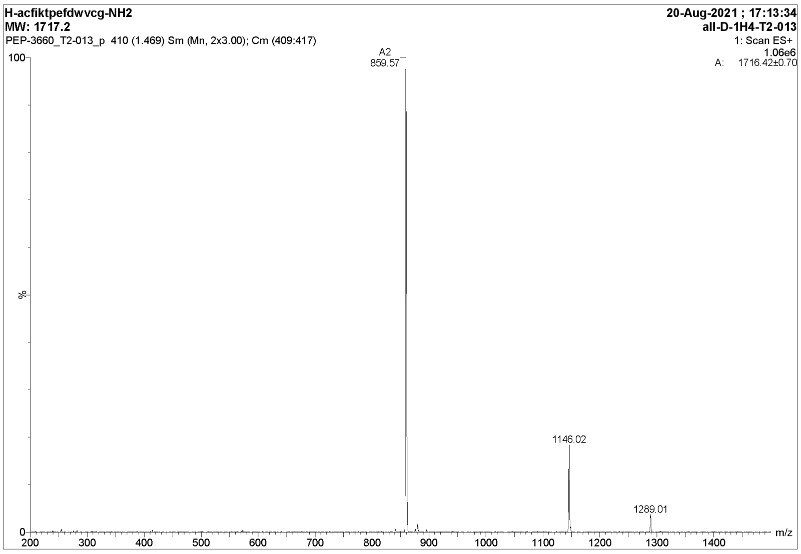

**Figure S12. LC-MS analysis of D-CLIPS$^{TM}$ peptide 1H4**

The HPLC trace (top) and ESI mass spectrum (bottom) of D-CLIPS$^{TM}$ peptide 1H4. Observed *m/z* signals: $[M+2H]^{2+}$ - 859.7, $[2M+3H]^{3+}$ - 1146.0, $[3M+4H]^{4+}$ - 1289.0; calculated *m/z* signals: $[M+2H]^{2+}$ - 859.6, $[2M+3H]^{3+}$ - 1145.8, $[3M+4H]^{4+}$ - 1288.9.

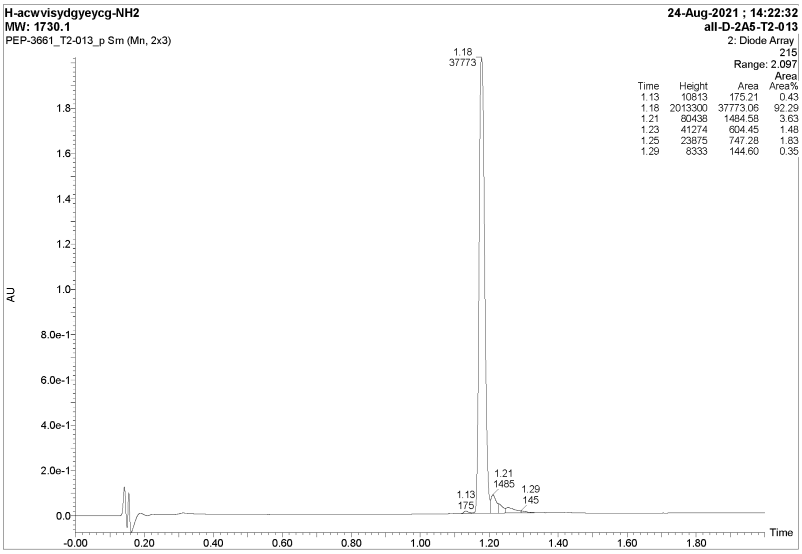

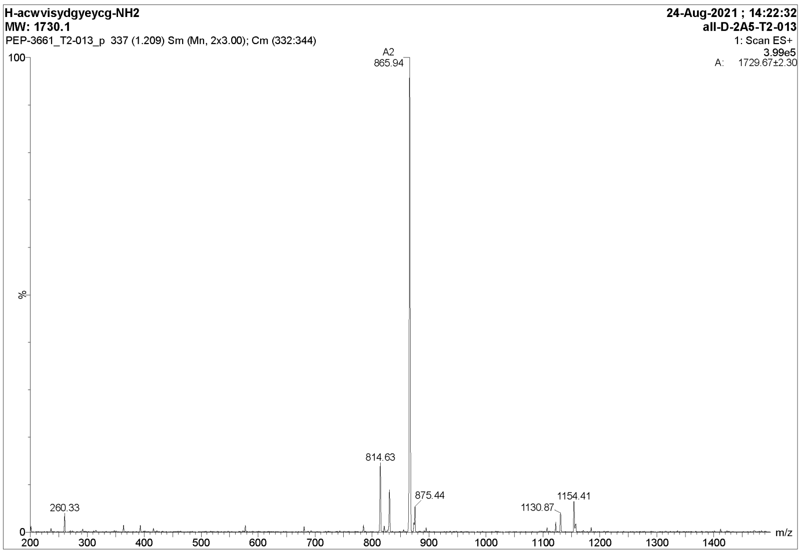

**Figure S13. LC-MS analysis of D-CLIPS$^{TM}$ peptide 2A5**

The HPLC trace (top) and ESI mass spectrum (bottom) of D-CLIPS$^{TM}$ peptide 2A5. Observed *m/z* signals: $[M+2H]^{2+}$ - 865.9, $[2M+3H]^{3+}$ - 1154.4, calculated *m/z* signals: $[M+2H]^{2+}$ - 866.1, $[2M+3H]^{3+}$ - 1154.4.

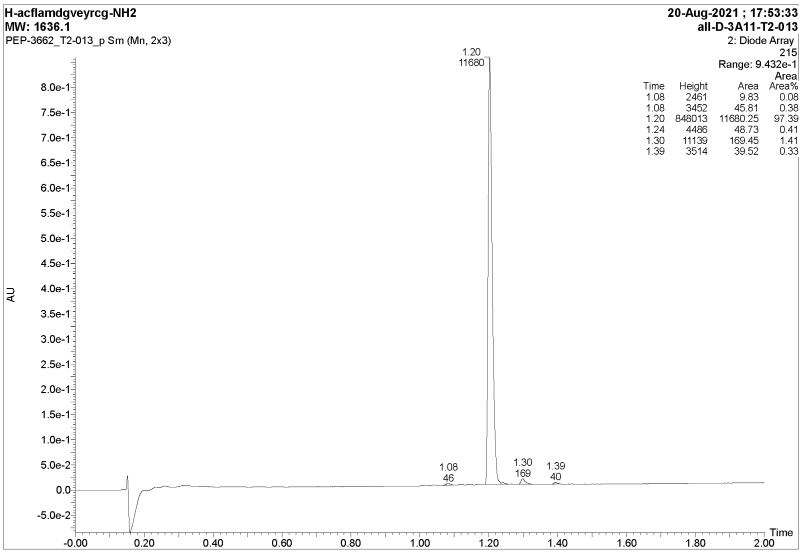

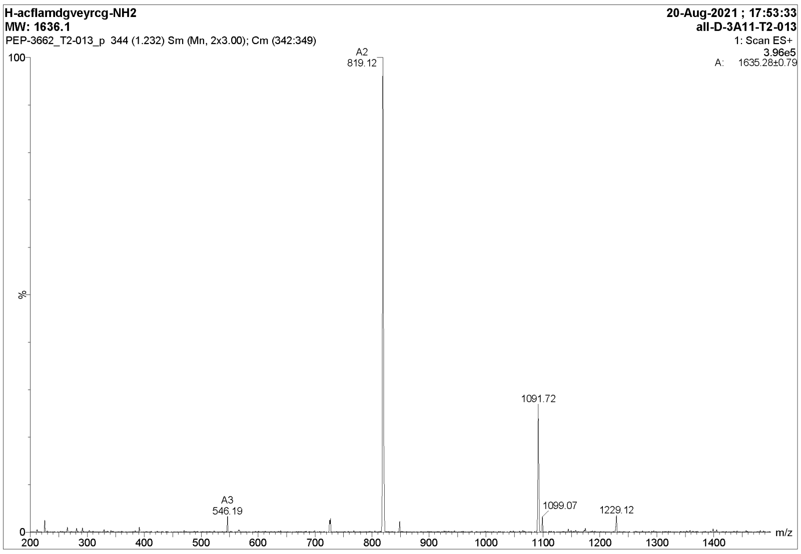

**Figure S14. LC-MS analysis of D-CLIPS™ peptide 3A11.**
The HPLC trace (top) and ESI mass spectrum (bottom) of D-CLIPS™ peptide 3A11. Observed *m/z* signals: $[M+2H]^{2+}$ - 819.1, $[2M+3H]^{3+}$ - 1091.7, $[3M+4H]^{4+}$ - 1229.1; calculated *m/z* signals: $[M+2H]^{2+}$ - 819.1, $[2M+3H]^{3+}$ - 1091.7, $[3M+4H]^{4+}$ - 1228.1.

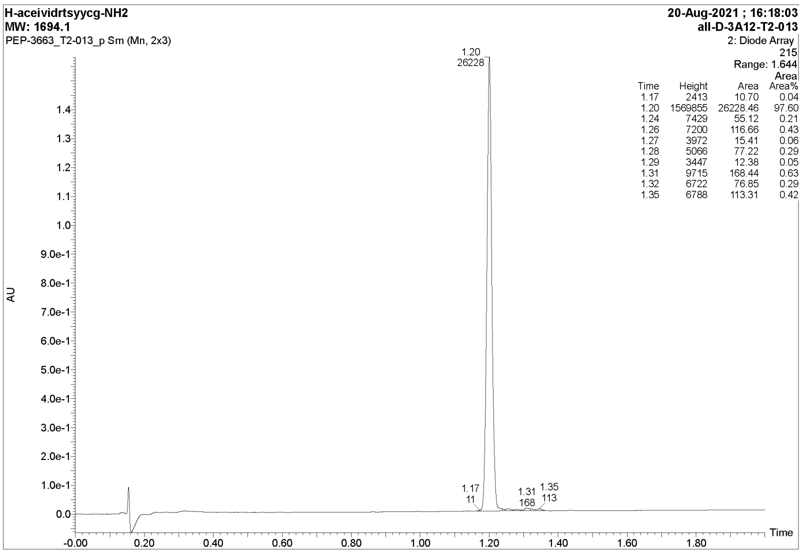

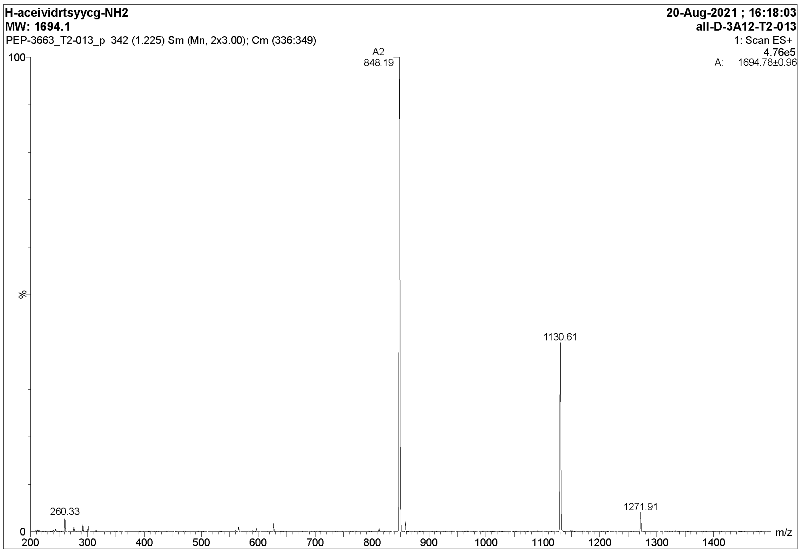

**Figure S15. LC-MS analysis of D-CLIPS$^{TM}$ peptide 3A12.**

The HPLC trace (top) and ESI mass spectrum (bottom) of D-CLIPS$^{TM}$ peptide 3A12 Observed *m/z* signals: $[M+2H]^{2+}$ - 848.2, $[2M+3H]^{3+}$ - 1130.6, $[3M+4H]^{4+}$ - 1271.9; calculated *m/z* signals: $[M+2H]^{2+}$ - 848.1, $[2M+3H]^{3+}$ - 1130.4, $[3M+4H]^{4+}$ - 1271.2.

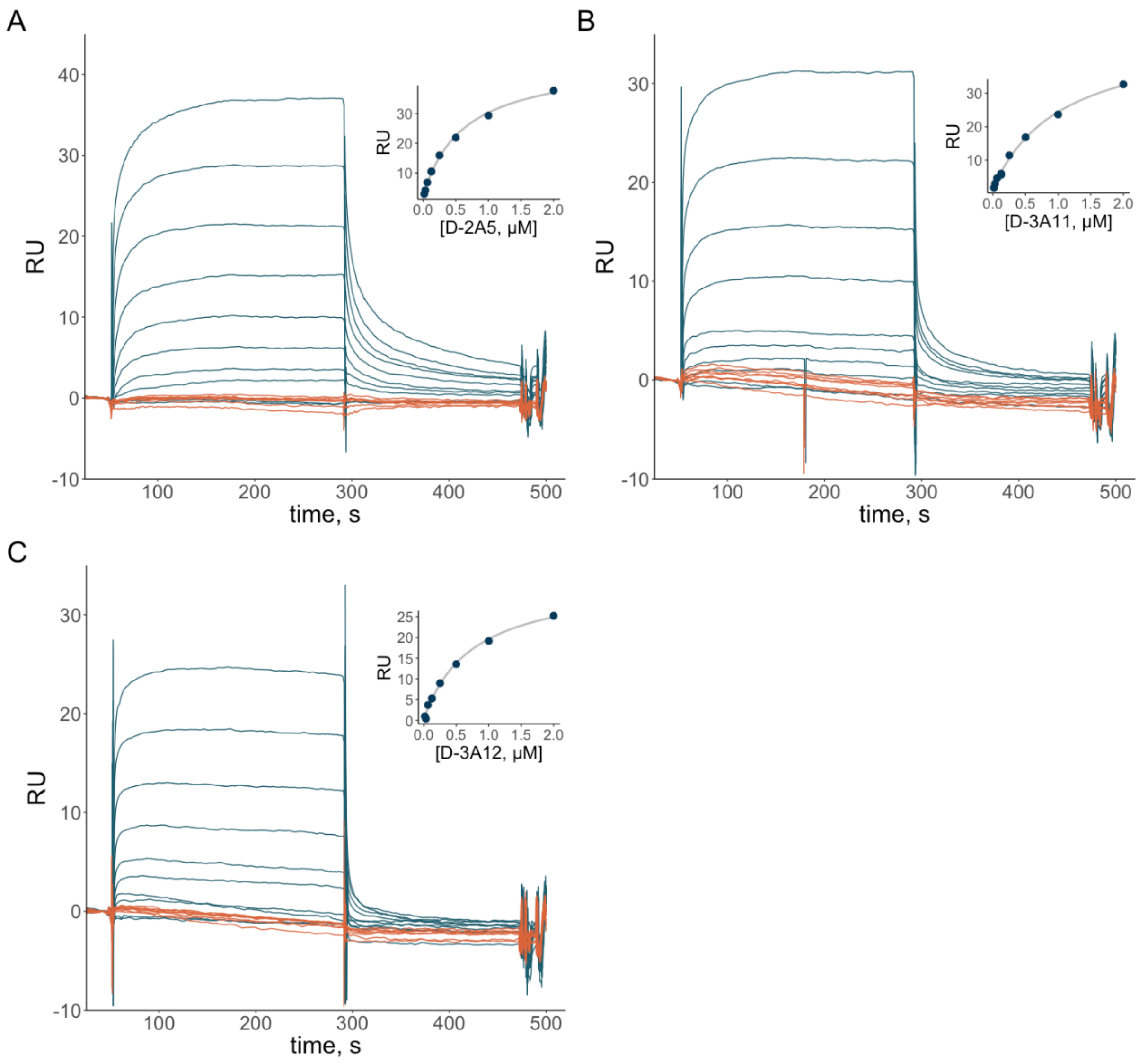

**Figure S16. Binding of D-peptides to L- and D-CXCL8.**
SPR biosensor analysis of interactions of D-CLIPS™ peptide 2A5 (A), D-CLIPS™ peptide 3A11 (B), and D-CLIPS™ peptide 3A12 (C) with D-CXCL8-KGG-biotin (orange) and L-CXCL8-KGG-biotin (blue) at concentrations 15 nM – 2 µM. Binding curves are shown in the inserts**.**

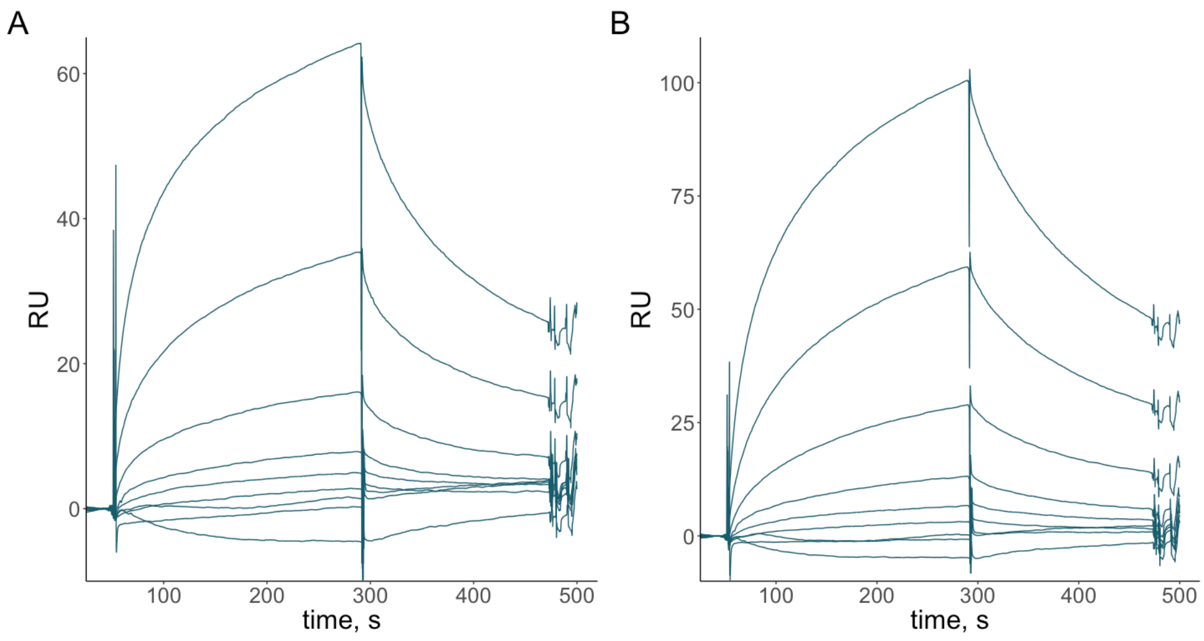

**Figure S17. Binding of D-2A5 to L-CCL5 and L-CXCL4**

SPR biosensor analysis of the interaction of D-2A5 at concentrations 15 nM – 2000 nM with L-CXCL4 (A) and L-CCL5 (B).

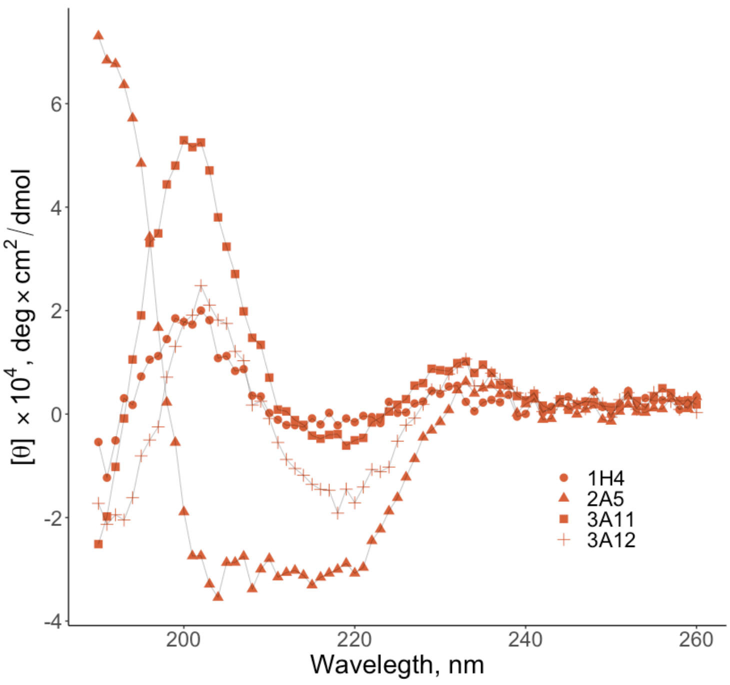

**Figure S18. Far-UV CD spectra of L-CLIPS peptides.** Spectra were recorded at a concentration of 0.1 mg/mL.

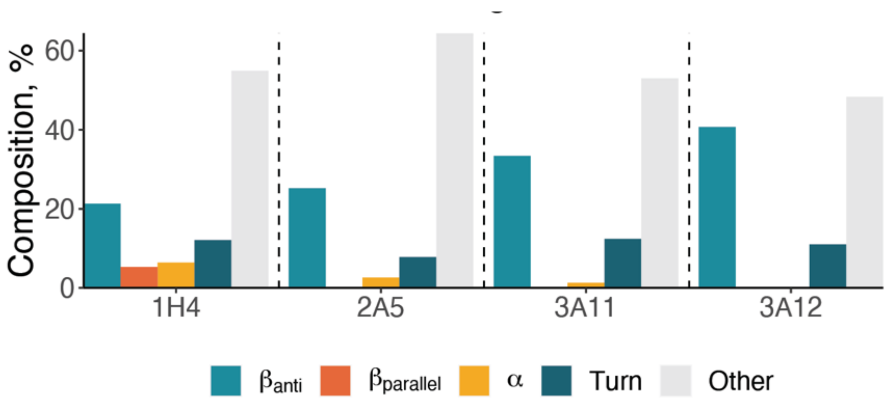

**Figure S19. Secondary structure composition of CLIPSTM peptides determined by the BeStSel algorithm using CD spectra at 0.1 mg/mL of L-peptides.**

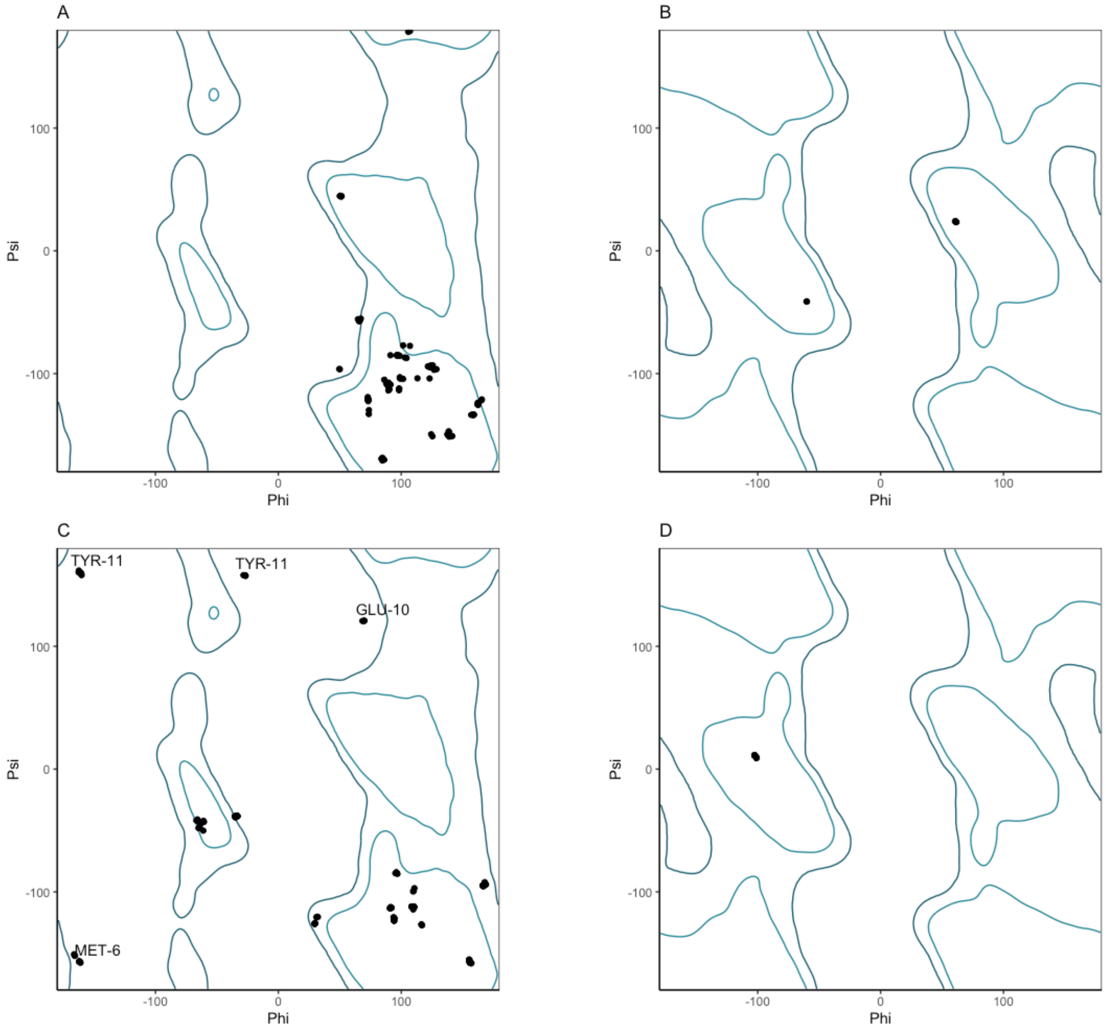

**Figure S20. Ramachandran plots of calculated D-peptide models**

Ramachandran plots for $\phi$ and $\psi$ angles for residues of 10 lowest-energy models of D-2A5 (A – general, B – Gly) and D-3A11 (C – general, D – Gly). Outliers in D-3A11 models are captioned.

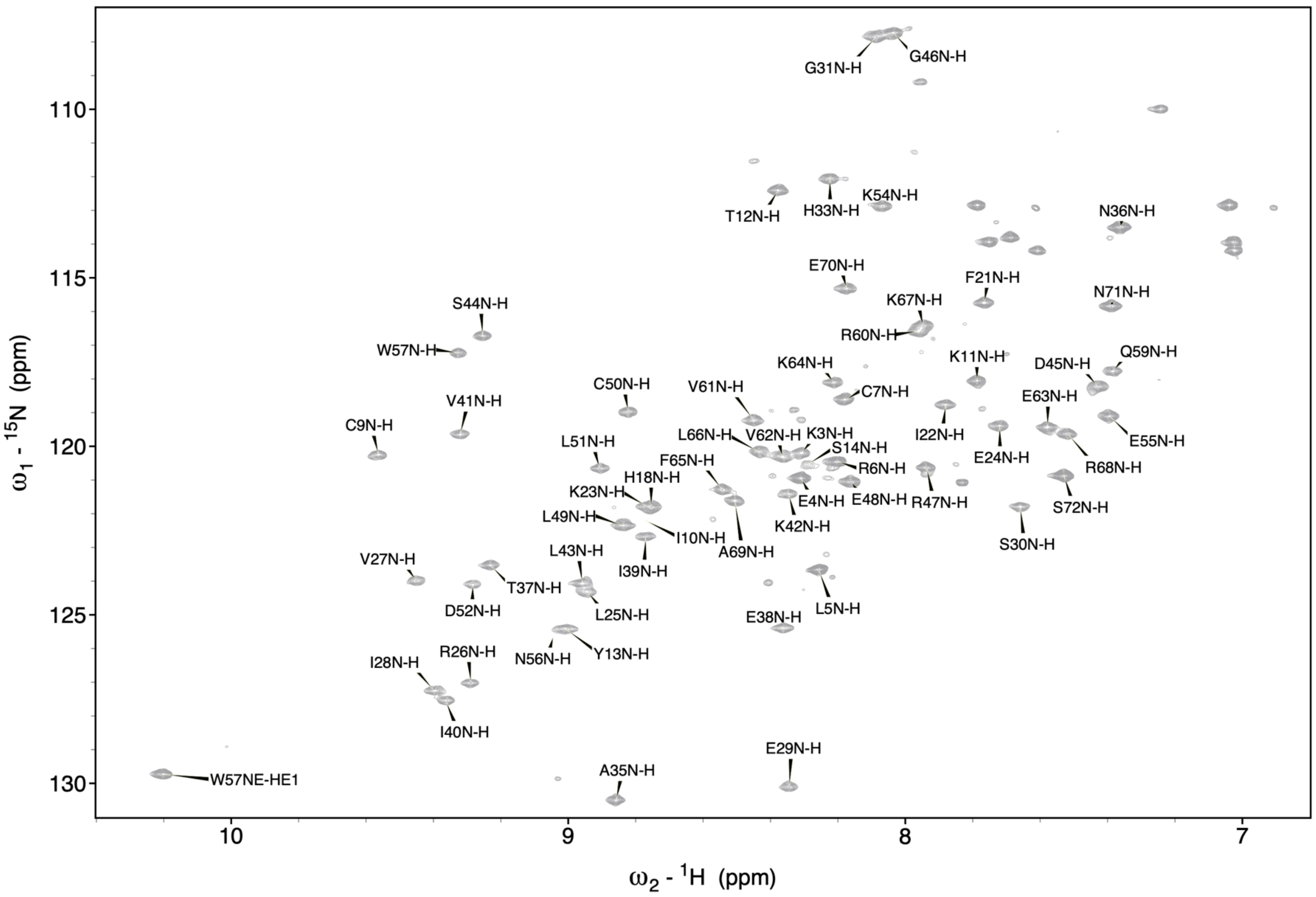

**Figure S21. $^{15}N$-$^{1}H$ HSQC spectra [$^{15}N$, $^{13}C$] L-CXCL8 dimer.**

Spectra of 25 μM [$^{15}N$, $^{13}C$] L-CXCL8 in 50 mM phosphate buffer, pH 7, 25 °C. Sidechain signals from Gln and Asn are hidden for visibility.

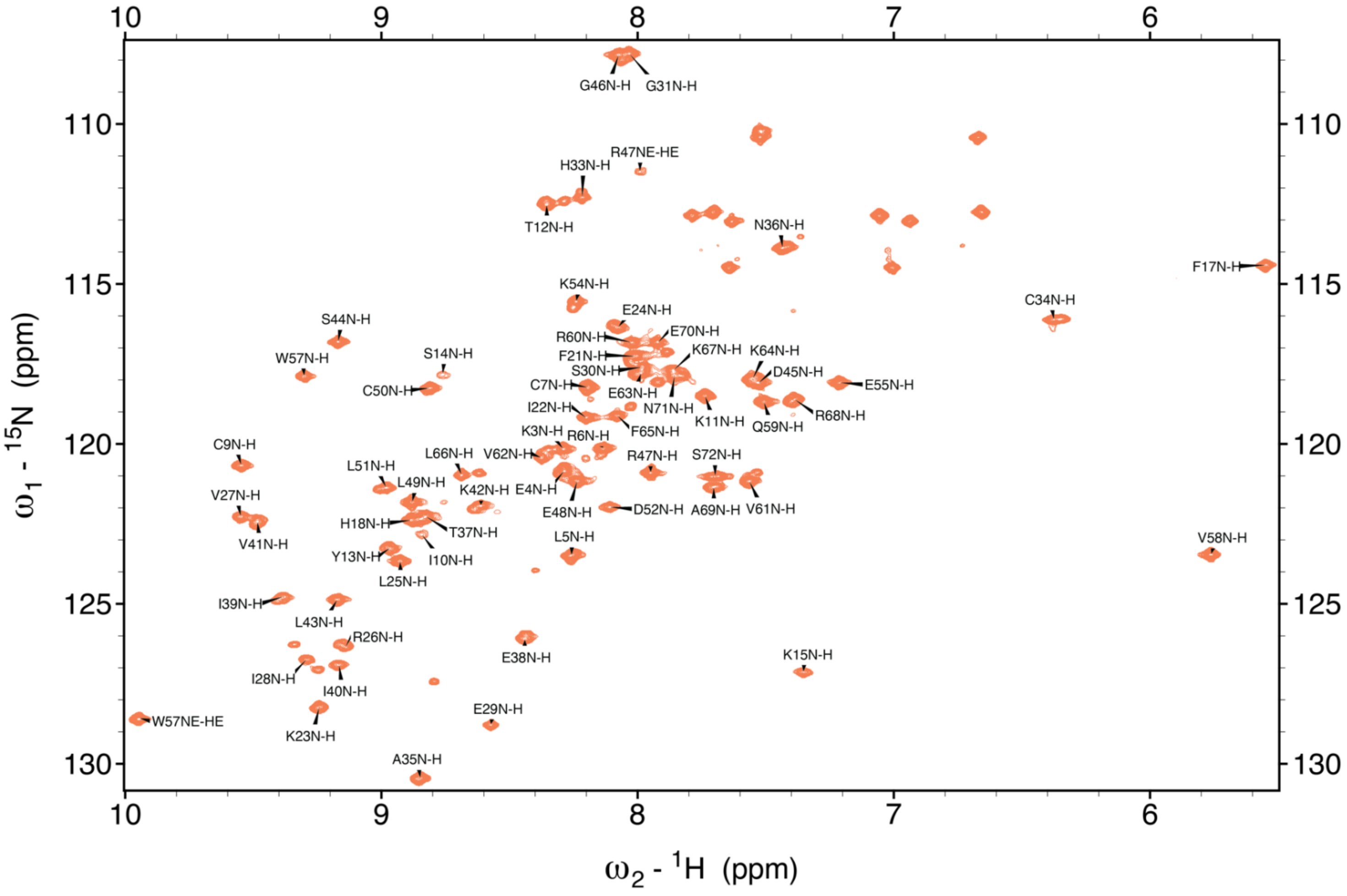

**Figure S22. $^{15}$N-$^{1}$H HSQC spectra [$^{15}$N, $^{13}$C] L-CXCL8/D-2A5 complex.**
Spectra of 59 µM [$^{15}$N, $^{13}$C] L-CXCL8 in the presence of 300 µM D-2A5 in 50 mM phosphate buffer, 10% DMSO, pH 7, 25 °C. Sidechain signals from Gln and Asn are hidden for visibility.

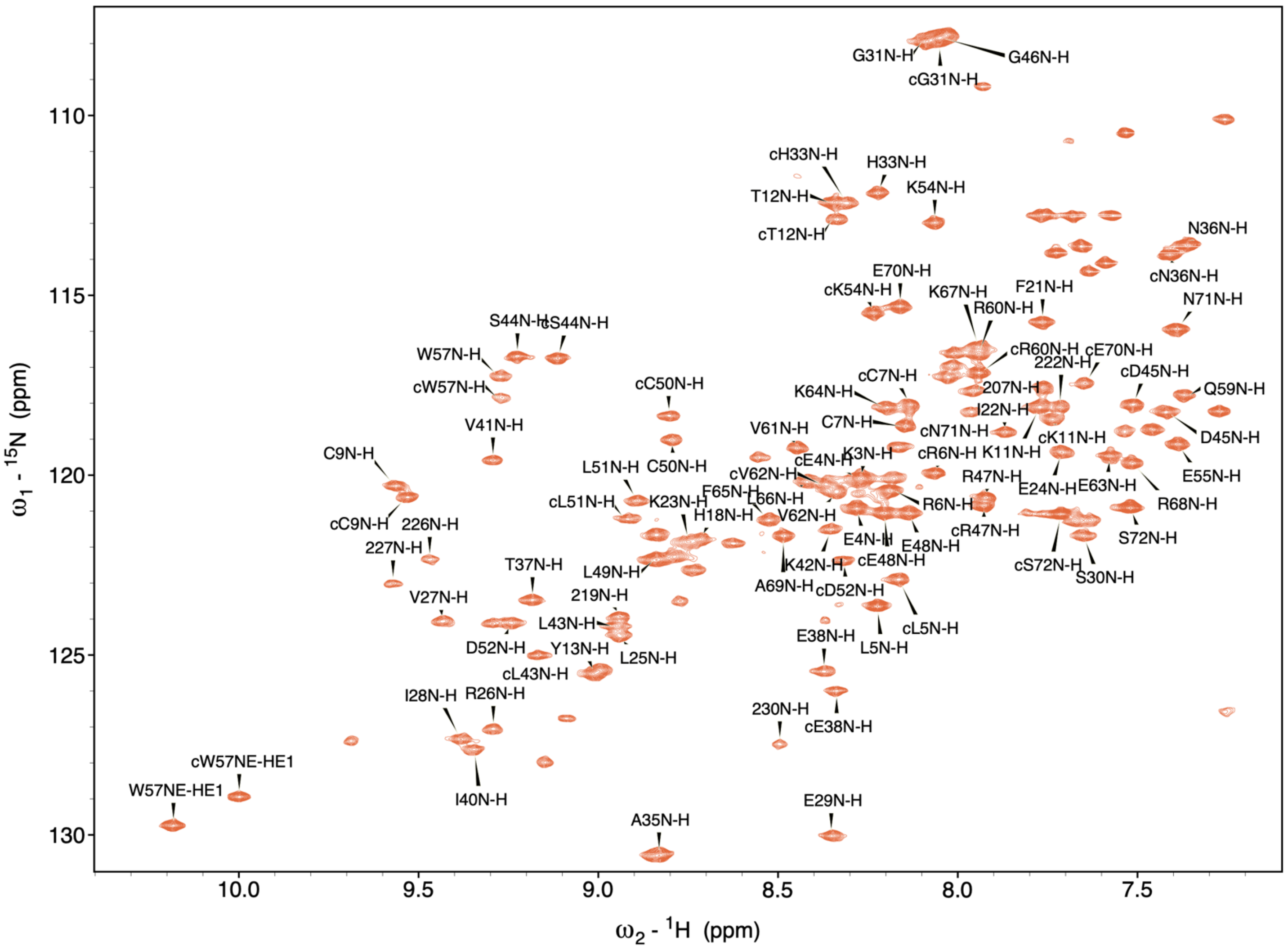

**Figure S23. $^{15}$N-$^{1}$H HSQC spectra of [$^{15}$N, 13C] L-CXCL8/D-3A11 complex**

The spectrum of 75 μM [$^{15}$N, $^{13}$C] L-CXCL8 in the presence of 300 μM D-3A11 in 50 mM phosphate buffer, pH 7, 25 °C. Unambiguously assigned signals from the bound L-CXCL8 are indicated with "c" and sidechain signals from Gln and Asn are hidden for visibility.
